# Solid-State Nanopore Real Time Assay for Monitoring Cas9 Endonuclease Reactivity

**DOI:** 10.1101/2024.09.20.612695

**Authors:** Chalmers C. Chau, Nicole E. Weckman, Emma E. Thomson, Paolo Actis

## Abstract

The field of nanopore sensing is now moving beyond nucleic acid sequencing. An exciting new avenue is the use of nanopore platforms for the monitoring of biochemical reactions. Biological nanopores have been used to investigate the dynamics of enzymatic reactions but solid-state nanopore approaches have lagged. This is partially due to the necessity of using higher salt conditions (*e.g.* 4M LiCl) to improve the signal-to-noise ratio. These conditions are often incompatible with monitoring biological activities such as restriction enzyme and CRISPR-Cas9 endonuclease activities, as their abilities to digest DNA are largely abolished under high salt conditions. We pioneered a polymer electrolyte based solid-state nanopore approach that can maintain a high signal-to-noise ratio even at a physiologically relevant salt concentration. Based on this, we report monitoring the endonuclease activities at physiological salt conditions in real time with a polymer electrolyte nanopore. We successfully monitor the dsDNA cleavage activity of the restriction enzyme SwaI in in a range of digestion buffers and the off-target activity of CRISPR-Cas9 RNP endonuclease in the presence of single base pair mismatches. This approach enables solid-state nanopore technology for the dynamic monitoring of biochemical reactions at single molecule resolution.

## Introduction

Single molecule analysis is advancing the understanding of fundamental biochemical and biophysical processes involving DNA, RNA, proteins, and cellular machinery ^1–3^. Numerous tools have been developed for single molecule analysis such as atomic force microscopy ^4^, super resolution fluorescent microscopy ^5^, optical tweezers ^6^, and single molecule Förster resonance energy transfer (sm-FRET) ^7^. Unlike many of these techniques that require fluorescent or chemical labelling of the analyte, nanopore sensors are a label-free single molecule analysis technique amenable to high throughput analyses ^3^. Nanopores were utilized for the sequencing of nucleic acids ^8^ but have also been extensively used for the biophysical characterization of a range of biomolecules ^9–13^. In a nanopore experiment, an analyte passes through the nanopore under the application of an electric field. This elicits a temporary modulation of the measured ion current through the pore, which depends on the volume and surface charge of the translocating analyte ^13^. Nanopores are most known for their commercial application for sequencing nucleic acids ^8^, however, they are broadly useful for the biophysical characterization of a range of biomolecules and their interactions^9–13^. Furthermore, nanopores provide real-time detection of analytes which makes them ideal as a technique to monitor reactions ^14,15^ or investigate biochemical processes such as monitoring enzymatic activity ^14–18^.

Enzymes such as endonucleases are biological catalysts of chemical reactions and play a major role in physiological processes and in synthetic biology ^19^, and biological nanopores have been used to investigate the dynamics of enzymatic reactions ^17,18,20–23^. The investigation of enzymatic reactions with biological pores often relies on immobilization of the enzyme at the nanopore or confinement of the enzyme in the pore ^14–16^. The size of solid-state nanopores can be easily tuned allowing for the direct study of both products of enzymatic reactions as well as the enzyme-substrate interactions without immobilization or confinement of the enzyme ^24–27^.

However, the use of solid-state nanopores to monitor enzymatic reactions has been limited, partially due to the need of using high electrolyte conditions (e.g. 4M LiCl) to improve the signal-to-noise ratio ^24–29^. As most enzymes are only catalytically active under specific salt concentration, these high salt conditions do not allow for the real time monitoring of enzymatic activities.

Here, we overcome this challenge by using a polymer electrolyte bath to enhance the solid-state nanopore sensing performance while still enabling measurements in physiologically relevant salt and buffer conditions ^30–34^. We demonstrate the capabilities of this system for monitoring real-time of the digestion activities of two different sequence specific endonucleases: restriction enzyme SwaI and CRISPR-Cas9. We develop and optimize the real-time quantitative analysis system using a well-understood restriction endonuclease, SwaI. We then demonstrate the proof-of-principle of real-time quantitative analysis of CRISPR-Cas9 on target and off target endonuclease activity, an understanding of which is critically important for applications in gene therapy or the wider biotechnology applications of CRISPR-CAS9 approaches.

With further development and optimisation, we envision that we can apply our solid-state nanopore kinetic measurement system beyond monitoring endonuclease activities, with many potential usages for monitoring assembly and disassembly of larger biological complexes. The tuneable pore size of our system opens the door to a broad range of applications studying biomolecules of various sizes, like facilitating the detection of the aggregation of protein aggregates ^30,35,36^.

## Results and discussion

### Real-time quantitative nanopore analysis system for endonuclease

We designed a 3kbp dsDNA (RS-dsDNA) with a restriction site for the restriction enzyme SwaI, at its centre ^37^, and designed the on and off target crRNA variants for the CRISPR-Cas9 digestion system to perform digestion at the centre of the RS-dsDNA ^38^. The successful digestion of the RS-dsDNA by either enzyme means the cleavage of the 3kbp RS-dsDNA, results in the production of two 1.5kbp dsDNA fragments. Our solid-state nanopore measurement system which can clearly distinguish the sizes of these dsDNA ^34^ based on peak amplitude of the translocation event alone and can monitor the population of these cleaved RS-dsDNA in real-time. For the SwaI endonuclease activity, we investigated the effect of the buffer composition with solid-state nanopore and showed that suboptimum buffer composition abolished enzyme activities, in agreement with gel electrophoresis data. For CRISPR-Cas9, our system allows us to study the effects of sequence mismatches between the crRNA and the target DNA sequence on the endonuclease activity. Mismatches led to slower cleavage activities and our system indicates that mismatches at position 1 and 4 upstream of the PAM site reduced the cleavage activities significantly.

We recently developed a new method to enhance the performance of a glass solid-state nanopore, by replacing the electrolyte bath (trans chamber) with a polymer electrolyte bath composed of 50% (w/v) poly(ethylene) glycol (PEG) and 0.1M KCl ^30–34^. With this method, the translocation of molecules from cis chamber (inside the nanopore) to trans chamber (outside the nanopore) generates a conductive pulse. Crucially, the sensitivity of the system only depends on the composition of the trans chamber bath while the composition of the solution within the cis chamber, where the analytes are placed, plays a negligible role in determining the signal-to-noise ratio ^31,33,34^. This allows the measurement to be operated at near or lower than physiological salt concentration without compromising the single-molecule sensitivity. The glass nanopore used here has a diameter of approximately 70nm (S.Figure 1) and its response is highly reproducible as demonstrated by I-V measurements (S.Figure 2), indicating high reproducibility on the geometry of the nanopore.

Through modern synthetic biochemistry and molecular cloning techniques ^39^ (**Error! Reference source not found.**A and S.Figure 3), we generated the 3kbp dsDNA (RS-dsDNA) containing a specific restriction site (5’-ATTT/AAAT-3’) at the centre of the RS-dsDNA which allows the restriction enzyme SwaI to digest it into 2 pieces of 1.5 kbp dsDNA ^37^, as shown in **Error! Reference source not found.**B.

We performed translocation of dsDNA of different sizes into the polymer electrolyte bath and observed that dsDNA of distinct sizes could be classified based on their peak amplitudes ^31,34^ (S.Figure 4A, refer to Supporting information note 1 for the mechanism explanation). We tested the translocation of the purified RS-dsDNA under different voltages (S.Figure 4B) and we selected −700 mV for the rest of the study, as it provided a suitable capture rate to facilitate statistical analyses. The concentration of the RS-dsDNA was determined to be optimal after screening different concentrations (S.Figure 4C). Next, we tested the ability of the system to distinguish the SwaI digested (1.5 kbp) and undigested RS-dsDNA (3.0 kbp), prior to experiment, the RS-dsDNA was digested with the restriction enzyme SwaI and the products were purified and diluted down to 10nM with 0.1M KCl. We observed translocation signals for both the RS-dsDNA and the digested RS-dsDNA (S.Figure 4D). The population scatter showed the major population shift from a higher current amplitude to a lower current amplitude after digestion, in line with our previous observations ^34^. The current peak amplitude of the single molecule translocation events where the undigested RS-dsDNA formed a population at about 0.3nA while the digested RS-dsDNA formed a population at about 0.2nA (S.Figure 4E), showing that the current peak amplitude component alone can be used to separate the population reliably, in contrast to dwell time component (S.Figure 4F). Due to the dimension of the nanopore used here (S.Figure 1), the translocation of the restriction enzyme could not be detected (S.Figure 5).

The detection of DNA translocating through a solid-state nanopore at high resolution has been reported extensively before, but most experiments are carried out at higher than physiological salt concentration (e.g. 0.5 to 4M) and with a monovalent salt like LiCl that is not commonly found in physiological conditions ^40–53^. However, performing single molecule detection with solid-state nanopores at high salt conditions can hinder or supress the activity of restriction enzymes. Most routinely used digestion buffers contain 50 to 100mM of NaCl (Table 1) and increasing the concentration of NaCl to 1M completely abolishes the SwaI activity (S.Figure 7). The high dependency on the salt concentration makes enzyme activities difficult to monitor using solid-state nanopores.

**Table 1.** Composition of digestion buffers.

| Buffer Name | Composition |
| --- | --- |
| Buffer 3.1 | 50mM Tris-HCl, 10mM MgCl<sub>2</sub>, 100mM NaCl, 100µg/ml BSA at pH 7.9 |
| Buffer 4 | 20mM Tris-OAc, 10mM MgOAc, 50mM KOAc, 1mM DTT, pH 7.9 |
| Buffer 2.1 | 50mM Tris-HCl, 10mM MgCl<sub>2</sub>, 50mM NaCl, 100µg/ml BSA, pH 7.9 |
| Cutsmart | 20mM Tris-OAc, 10mM MgOAc, 50mM KOAc, 100µg/ml BSA, pH 7.9 |

To overcome this challenge with traditional solid-state nanopore sensing, we leverage the benefits of the polymer electrolyte and developed an assay to monitor the dsDNA cleavage activities of the SwaI enzyme over time. This is done by monitoring the gradual reduction in the number of RS-dsDNA translocation events over time as it gets cleaved by SwaI. To prevent potential clogging of the dsDNA at the nanopore over time, we applied a waveform composed of 3.5 seconds of +100mV followed by 6 seconds of −700mV (S.Figure 8). This waveform allows us to control the delivery of the dsDNA on demand as the dsDNA will migrate away from the nanopore due to the voltage polarity ^54^. This waveform is looped for 180 to 360 times (30 min to 1 hour), allowing the real time monitoring of the enzymatic reaction.

We first demonstrated that the digestion reaction carried out inside the glass nanopore capillary provides comparable results to the reaction performed in a standard polypropylene reaction tube (S.Figure 9). Next, the mixture composed of 1× buffer 3.1 (Table 1), 10nM RS-dsDNA and 5 units of SwaI was prepared and immediately loaded into the cis chamber of the glass nanopore, which was then immersed in the polymer electrolyte bath followed by the application of the waveform described above (Figure 2A). The translocation events at 1, 10, 20 and finally at 30 min were isolated and 20 random events were overlapped as shown in Figure 2B, to show the visual progression of the reduction in the current peak amplitudes at different time points. Similarly, Figure 2C showed the population distribution of the translocation events at 1, 10, 20 and 30 min. The higher peak amplitude population was the major population at the 1 min time point, indicating the presence of the RS-dsDNA, this population slowly reduced at 10 min, 20 min and finally only a minor population of the RS-dsDNA could be detected at 30 min.

**Figure 1.** The generation of the restriction site containing 3 kbp dsDNA. **(A)**, Schematic illustration of the generation of the restriction site containing 3 kbp dsDNA (RS-dsDNA). **(B)**, Agarose gel electrophoresis analysis of the undigested RS-dsDNA and the digested RS-dsDNA, the 3 kbp original fragment was digested into 1.5 kbp dsDNA.

**Figure 2.**
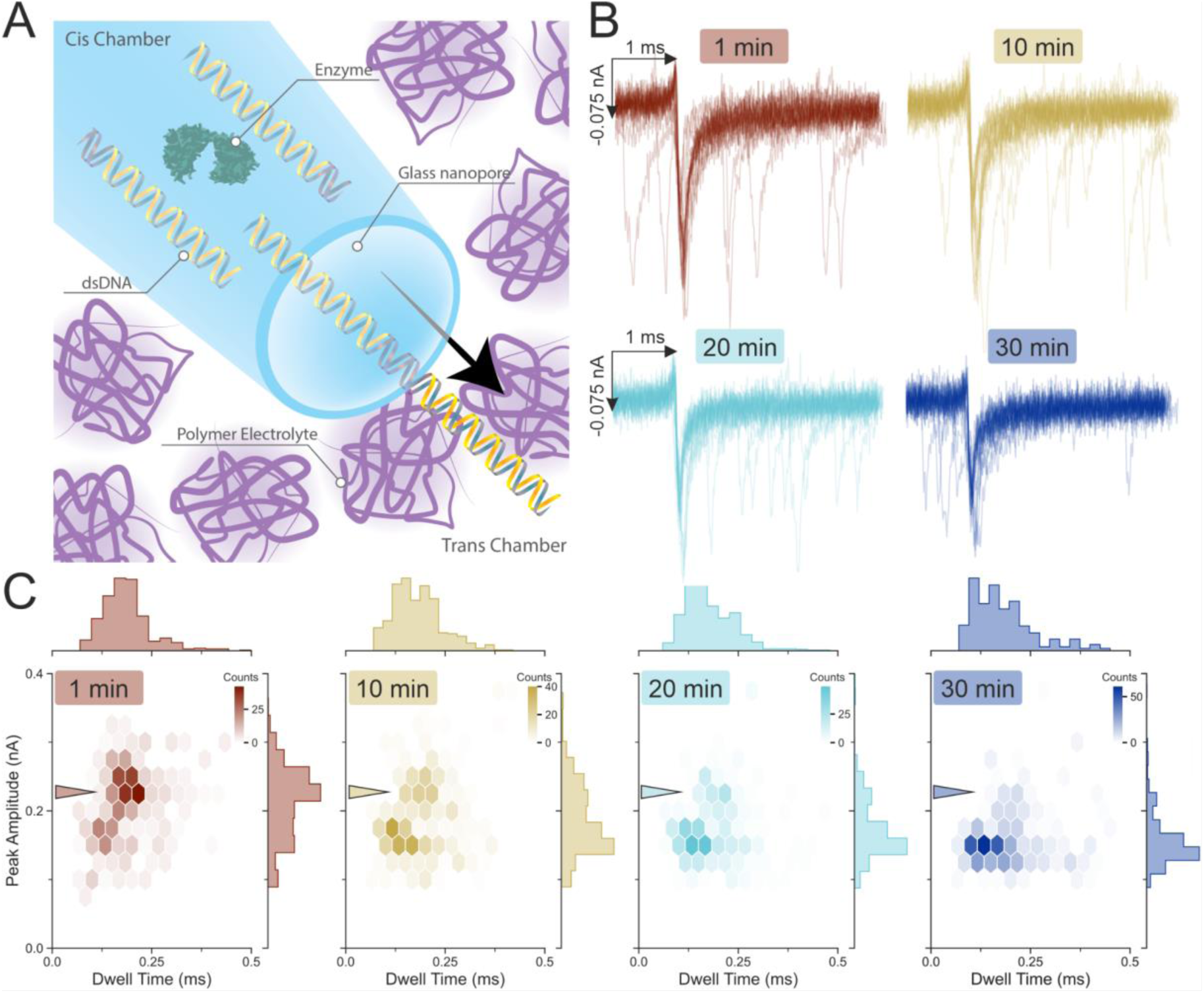
Restriction enzyme SwaI cleavage activities. **(A),** Schematic illustration of the glass nanopore detection set-up. The cis chamber of the glass nanopore is filled with the restriction enzyme SwaI, the RS-dsDNA, the trans chamber is composed of a polymer electrolyte mixture (0.1M KCl, 50% (w/v) PEG 35K), negative voltage caused the dsDNA to migrate from cis to trans chamber. The RS-dsDNA was mixed with the SwaI restriction enzyme and digestion buffer to a final concentration of 9.24 nM RS-dsDNA, 5 units of enzyme and 1× digestion buffer**. (B)**, Random 20 translocation event peaks plotted as overlay. The cis chamber of the glass nanopore was filled with 10nM RS-dsDNA diluted with buffer 3.1 (Table 1) containing 5 units of SwaI enzyme. **(C),** The population distribution of the translocation events at 1, 10, 20 and 30 min. The arrowheads across the four plots point at the population centred at approximately 0.2 ms and 0.25 nA. This population is attributed to the larger RS-dsDNA prior to digestion. As the time progress to 10, 20 and finally at 30 min, this population gradually disappears and a secondary population centred at approximately 0.15 ms and 0.15 nA begins to emerge, the side histograms of the peak amplitude axes show the emergence of the 0.15 nA population. Colour bar represents the count of events found in each hexagon.

The dynamically shifting population could be difficult to capture through scatter plots, instead, the ridgeline plot was used to visualise the population changes over time. The ridgeline plots were composed of multiple non-parametric kernel density estimation (KDE) of the probability density functions (PDF) of the peak amplitude component across the observation time every min (Figure 3A). The higher peak amplitude population centred at around 0.25 nA could be seen slowly reduced over the course of 1 hour, while lower peak amplitudes centred at around 0.15nA started to emerge and became the major population after around 8 min. This observation agreed with the gel analysis on the digestion kinetics of the enzyme (S.Figure 10) indicating that the nanoconfinement of the SwaI within the nanopore and capillary did not alter its dsDNA cleavage activities.

**Figure 3.**
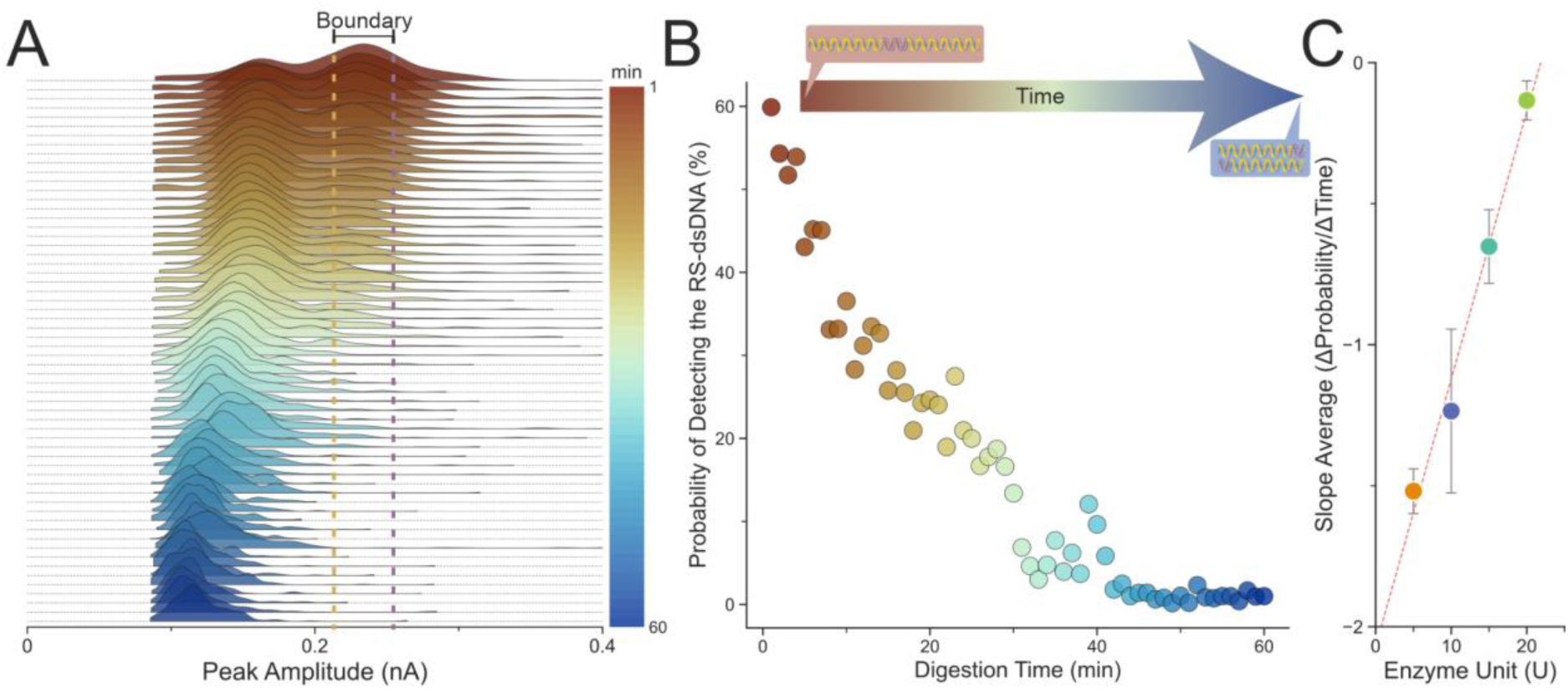
Digestion of RS-dsDNA monitored over the course of one hour. The RS-dsDNA was mixed with the SwaI restriction enzyme and digestion buffer to a final concentration of 9.24 nM RS-dsDNA, 5 units of enzyme and 1× digestion buffer. **(A)**, Ridgeline plot shows the gradual population changes from 1 min to 60 min. **(B)**, The probability of detecting the translocation of RS-dsDNA drops from 60% to near 0%. Two boundaries were defined as the ±10% of the peak value of the RS-dsDNA using the 1-min data, the same boundaries were applied across all the data. The probability value was calculated by integrating the area under the curves (AUC) between the boundaries shown in panel (A), the initial starting percentage changes according to the width of the boundaries. **(C)**, The enzyme reaction rate (slope average) plotted against the concentration of enzyme. The enzyme reaction rate was calculated by fitting a linear regression line at the first 15 mins (initial velocity region) of digestion under different enzyme concentrations (n=3), the reaction rate obtained from the linear regression line is thus defined as the changes in probability of detecting the RS-dsDNA over the changes in time. The coefficient of determination for the fit is R^2^ = 0.9822. Error bars represent standard error of the mean between the slope value between measurements. According to the manufacturer, A single unit of SwaI is defined as the amount of enzyme required to digest 1 µg of pXba DNA in 1 hour at 25°C in a total reaction volume of 50 µl.

To quantify the population differences over time, we utilised the fundamental properties of PDF (regardless of parametric and non-parametric derived PDF). The KDE estimates the PDF of the population distribution, the area under the curve (AUC) of the PDF must sum to 1 as all samples must fall within this PDF, the probability of an event to be bound within this PDF is 100%.

Subsequently, the AUC bound by two boundaries (Figure 3A) at the horizontal axis of the PDF will return the probability of the population, in this case, the probability of detecting a single molecule event with a current amplitude that falls within that AUC. Two boundaries (upper and lower) were defined by ±10% of the peak value of the RS-dsDNA population from the 1 min trace (Supporting Method for detail explanations on the boundaries selection and probability calculation method), 1 min trace was selected as for most samples the 1 min trace would contain both the RS-dsDNA and 1.5 kbp dsDNA populations. These boundaries were fixed and applied across all PDFs within the observation time, and the probabilities were calculated for each min. This analytical pipeline was used to quantify the variation of the dsDNA populations (3kbp vs 1.5kbp) over time. Figure 3B showed a scatter plot of the probability of detecting the RS-dsDNA translocation as a function of the digestion time. The probability dropped from close to 60% to around 15% after 30 min of digestion, reaching near 0% at around 40 min suggesting full digestion of the RS-dsDNA by SwaI. Additional SwaI concentrations ranging from 5U to 20U were monitored and their rates of the reactions were calculated. Through plotting the rate of reaction against the SwaI concentration, our data showed that the SwaI’s DNA cleavage activity is linearly proportional to increasing concentration of SwaI (Figure 3C).

### Effects of Salt on SwaI cleavage activity

The recommended buffer composition for the SwaI (Buffer 3.1) and its components are presented in Table 1. Alternative buffers can also be used (such as the Buffer 4, Buffer 2.1 and Cutsmart), however, the activities of SwaI will be reduced, as mentioned by the supplier and also tested (Figure 4A, B). Buffer 2.1 and Buffer 3.1 share similarities in the compositions of the buffer except that Buffer 2.1 has lower NaCl concentration (50mM NaCl) compared to Buffer 3.1 (100mM NaCl), while in Buffer 4 and Cutsmart, the NaCl is replaced with KOAc. These subtle differences in the composition of the buffer greatly affect the activity of SwaI.

**Figure 4.**
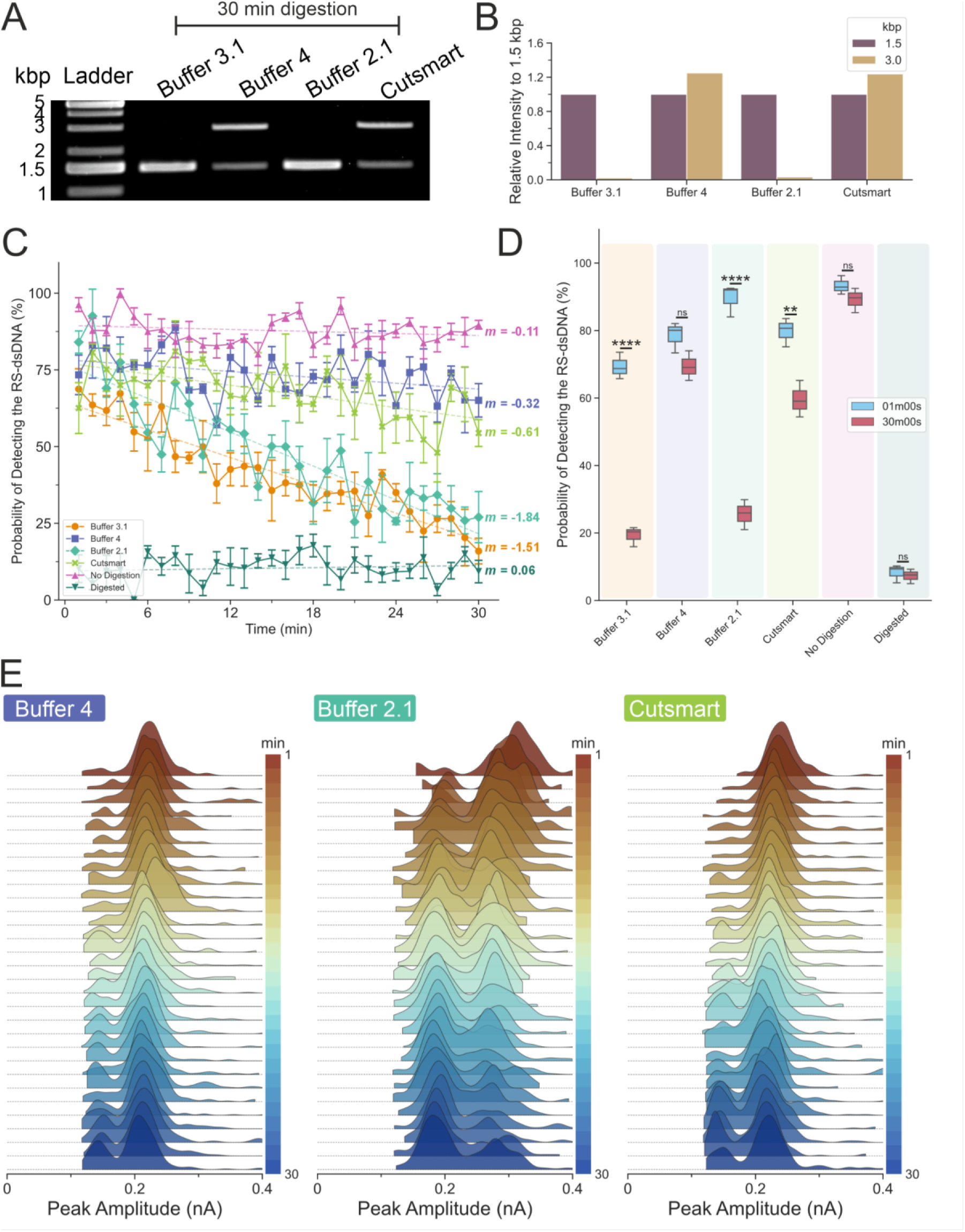
Buffer dependent restriction enzyme kinetics. **(A),** Restriction digestion of the RS-dsDNA in different buffer. The optimal buffer for the restriction enzyme SwaI is the buffer 3.1, as recommended by the supplier. Three other buffers (4, 2.1 and cutsmart) were tested, and the activity of SwaI varied and resulted in lower digestion activity in buffer 4 and cutsmart. **(B)**, The gel band intensity was quantified and calculated relative to the 1.5 kbp’s band intensity within the sample lane (self-reference). **(C),** Probability of detecting 3kbp dsDNA as a function of the digestion time for all the buffer tested and controls. *m* is the slope after fitting with the linear fit to the experimental data. Error bars are standard error of the mean. **(D),** Box plot comparing the probability of detecting the RS-dsDNA at 1 min and at 30 min. Two-tailed unpaired T test was used to test the differences between the distribution of the probabilities to detect RS-dsDNA at 1 min and at 30 min, there are significant differences for buffer 3.1, buffer 2.1 and cutsmart at 1 min and at 30 min. The calculated values for buffer 3.1 at 1 min is 69.35±3.94% and 19.28±2.95% at 30 min respectively, for buffer 4 at 1 min is 78.51±4.55% and 69.43±4.4% respectively, for buffer 2.1 at 1 min is 89.54±4.77% and 25.6±4.44% respectively, for cutsmart at 1 min is 79.81±4.23% and 59.56±5.42% respectively, for no digestion at 1 min is 93.31±2.76% and 89.17±3.62% respectively, for digested at 1 min is 8.52±1.72% and 7.26±2.15% respectively. (ns, not significant; ****, P ˂0.0001; **, P˂0.005; data assume normal distribution; Levene’s test (P>0.05) indicates data are homoscedasticity; N=3). **(E),** Ridgeline plots for buffer 4, buffer 2.1 and Cutsmart. The cis chamber of the glass nanopore was filled with 10nM RS-dsDNA diluted with either Buffer 3.1, Buffer 4, Buffer 2.1 or Cutsmart (Table 1) containing 5 units of SwaI enzyme.

As our nanopore set-up was inert to buffer composition in the cis chamber ^31,34^. This offered an opportunity for us to monitor the activities of SwaI under a slightly modified buffer composition in real-time. We replaced the buffer in the final reaction mixture prior to loading into the glass nanopore and monitored the cleavage activity of the SwaI enzyme under different buffer conditions (Figure 4E). Two control experiments were also performed where one contained the RS-dsDNA without the addition of SwaI enzyme (S.Figure 11), and the other where the RS-dsDNA were digested for 30 min before loading into the glass nanopore (S.Figure 12).

The probabilities of detecting the translocation of RS-dsDNA over 30 min of digestion time for different buffers were plotted in Figure 4C. The no enzyme controls showed consistently high probability (near 75%) of detecting the RS-dsDNA. The values did not change over time as indicated by the slope value of −0.03, indicating that, as expected, the materials inside the nanopore did not change. Similarly, the completely digested RS-dsDNA showed a low probability (near 12.5%) of detecting the ds-dsDNA, and this value did not change over the course of the digestion time (slope value = −0.03). When comparing the probabilities at 1 min and 30 min, there were no significant differences (Figure 4D).

In agreement with the gel electrophoresis data (Figure 4A, B), the enzyme retained its activity in both Buffer 3.1 and Buffer 2.1. The probability of detecting the RS-dsDNA dropped to near 25% after 30 min from an initial value higher than 50%. Their slope values were −1.15 and −1.41 respectively, suggesting a rapid decline in the number of RS-dsDNA available in the buffer over a short period of time, as also evidenced by comparing the probabilities at 1 min and 30 min (Figure 4D). Lastly, the digestion carried out with the non-ideal buffers (Buffer 4 and Cutsmart), both produced small changes in the probabilities over time (slope value = −0.24 and −0.46 respectively), indicating that while the enzyme was not as active as in Buffer 3.1 and Buffer 2.1, the enzyme could still digest the RS-dsDNA, but at a slower rate. Statistic testing indicated a significant difference between the 1 min and 30 min for the Cutsmart buffer, indicating that SwaI with Cutsmart buffer potentially operates at a slightly faster rate than in Buffer 4. These observations overall agreed with the gel electrophoresis data where both Buffer 4 and Cutsmart showed reduced activity when compared to Buffer 3.1 and Buffer 2.1 (Figure 4A).

### Effects of RNA:DNA mismatches on CRISPR-Cas9 cleavage activity

Part of the prokaryotic adaptive immunity mechanism used to cleave invading nucleic acids ^38^, the CRISPR-Cas9 is a unique, RNA-guided endonuclease; its dsDNA cleavage activity relies on a Cas9 ribonucleoprotein (RNP) complex composed of a Cas9 protein, a tracrRNA and a crRNA ^55–58^. The Cas9 RNP scans the target DNA to look for a short trinucleotide site – a, protospacer adjacent motif (PAM), site – and once a PAM site is identified, the target DNA sequence upstream of the PAM site is checked for complementarity against the crRNA. If complementary, a crRNA:DNA heteroduplex is formed and triggers the conformational activation of the Cas9 RNP’s HNH endonuclease domain to cleave the target DNA strand 3-4 nucleotides upstream of the PAM sequence ^55–58^. Notably, the Cas9 RNP does not dissociate from the DNA after cleavage ^59^ unlike restriction enzymes where they can proceed to cleave the next molecule of DNA.

The crRNA sequence can be designed to be complementary to different DNA sequences; targeting specific sequences in the genome enables the possibility of targeted gene editing. Thus Cas9 is widely studied for its applications in genome engineering and therapeutic potential for correcting genetic disorders ^60^. However, one challenge with using Cas9 RNP for genome editing, is that it shows off target activity. This happens when the crRNA is not fully complementary to the target DNA sequence but is still able to form a heteroduplex and triggers the HNH endonuclease domain ^61,62^. Depending on the position of the mismatch between the crRNA and the target DNA, the cleavage activity can be slower, or abolished ^63–67^.

The impact of mismatches and off target effects are critical problems to understand and address to unlock the potential of Cas9 for genome engineering. Traditional methods used to monitor the Cas9 RNP activity typically rely on sequencing ^63,67^ and electrophoresis ^64^, which can be costly and slow and often lack real-time kinetic information. Solid-state nanopores have been used to study the binding affinities between the Cas9 RNP and the target DNA through the binding of catalytically inactive dCas9 ^25–27^, including studying the effects of mismatches on the binding of dCas9 ^26^. However, the high salt conditions used in these studies can interact with the dsDNA backbone ^45,68,69^, leading to overwind in the dsDNA ^70,71^. The Cas9 RNP cleavage activity depends on unwinding the DNA duplex to form the heteroduplex and trigger the HNH endonuclease domain ^58,72,73^. Thus, increasing overwinding of dsDNA at high salt conditions is associated with reduced activity of the Cas9 RNP ^72^.

Our polymer electrolyte nanopore sensing system is ideal for rapidly studying the impact of mismatches on Cas9 RNP activity in real-time in physiologically relevant conditions, we identified a PAM site near the SwaI restriction site of the RS-dsDNA and designed a crRNA sequence to target the DNA 30 bp downstream of the site. Similar to SwaI, the DNA cleavage by the Cas9 RNP caused the RS-dsDNA to be separated into two 1.5 kbp dsDNA (Figure 5A, S.Figure 13). Additionally, we generated 4 off target crRNA variants with mismatches at different positions (Figure 5B). The mismatches introduced are rU-dG, rG-dT and rU-dC at position 1, 2, 3 or 4, upstream of the PAM site. The four off target crRNA variants and the on-target crRNA were assembled into five different Cas9 RNPs which were used to digest the RS-dsDNA. These were first assayed using traditional agarose gel electrophoresis with a single time point measurement after incubation at 25°C (room temperature) for 30 min, and then overnight at 4°C (Figure 5C). Within 30 min of incubation at room temperature the on-target crRNA led to the formation of the 1.5 kbp fragments. Similarly, the rG-dT variants at position 2 and 3 upstream of PAM site also successfully cleaved into 1.5 kbp fragments. In contrast, the rU-dG variant at position 1 and rU-dC variant at position 4 resulted in incomplete digestions by 30 min. After overnight incubation of the mixture, additional 1.5 kbp dsDNA fragments were formed with the off target 1 and 4 variants but the digestions were still incomplete.

**Figure 5.**
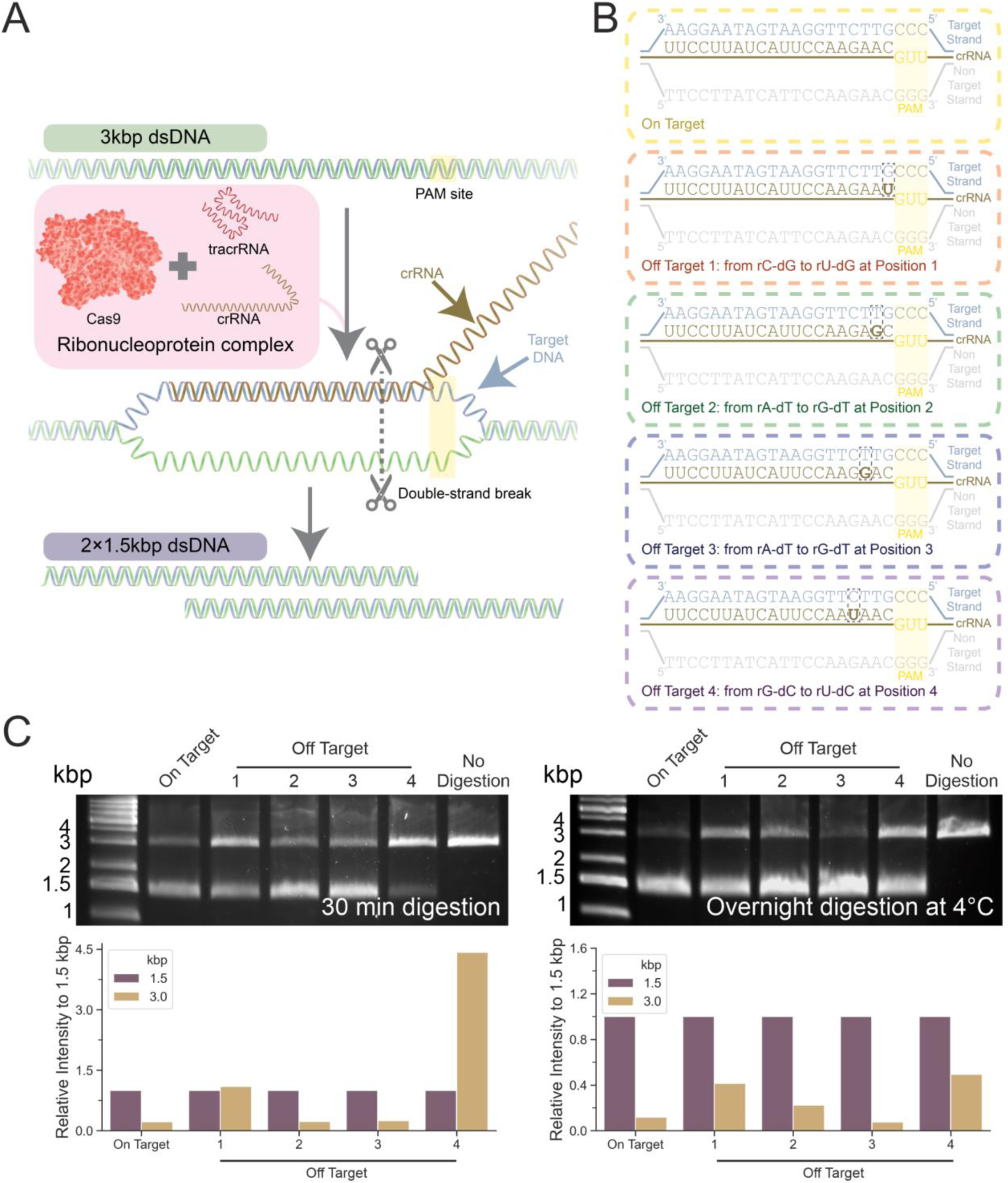
CRISPR-Cas9 mediated dsDNA cleavage. **(A),** Schematic illustrating the process of the Cas9 mediated cleavage on the 3kbp RS-dsDNA. The Cas9 was mixed with tracrRNA and crRNA to form the ribonucleoprotein (RNP) complex. The RNA guides the Cas9 RNP to the position next to PAM site, the Cas9 RNP then carries out double stranded break (DSB) 3-4 bp upstream of the PAM site to cleave the DNA. This resulted in the formation of 2×1.5 kbp dsDNA. **(B),** On target and off target crRNA sequence. The full complementary sequence (On target) and the off target variations at different positions upstream of the PAM site and different mismatches. **(C),** Agarose gel electrophoresis at 30 min incubation at 25°C and overnight incubation at 4°C, Cas9 RNPs were formed with the on target crRNA or off target crRNA variants. The gel band intensity was quantified and calculated relative to the 1.5 kbp’s band intensity within the sample lane (self-reference).

We then used our polymer electrolyte nanopore sensor to monitor the Cas9 RNP’s cleavage activity in real-time at a physiological relevant salt condition of 111 mM NaCl. We monitored the digestion of RS-dsDNA by different variants of the Cas9 RNPs over 30 min (Figure 6A), the off target 1 and 4 variants showed little changes in the population of the 3.0 kbp dsDNA (peak amplitude of ∼0.3 nA), the off target 2 and 3 variants showed a gradual reduction of 3.0 kbp population, indicated the RS-dsDNA was getting digested inside the nanopore, despite the presence of the mismatches. However, this was occurring at a much slower rate than the on target Cas9 RNP digestion. For the on-target digestion, most of the population that could be detected were 1.5 kbp fragments from the beginning of the experiment, indicating the RS-dsDNA were nearly fully digested by the time the assay started. Indeed, studies ^63,73^ showed that on target cleavage could cleave 80% of materials within 3 to 40 seconds and our method had a minimum delay of 60 seconds from mixing to measurement.

**Figure 6.**
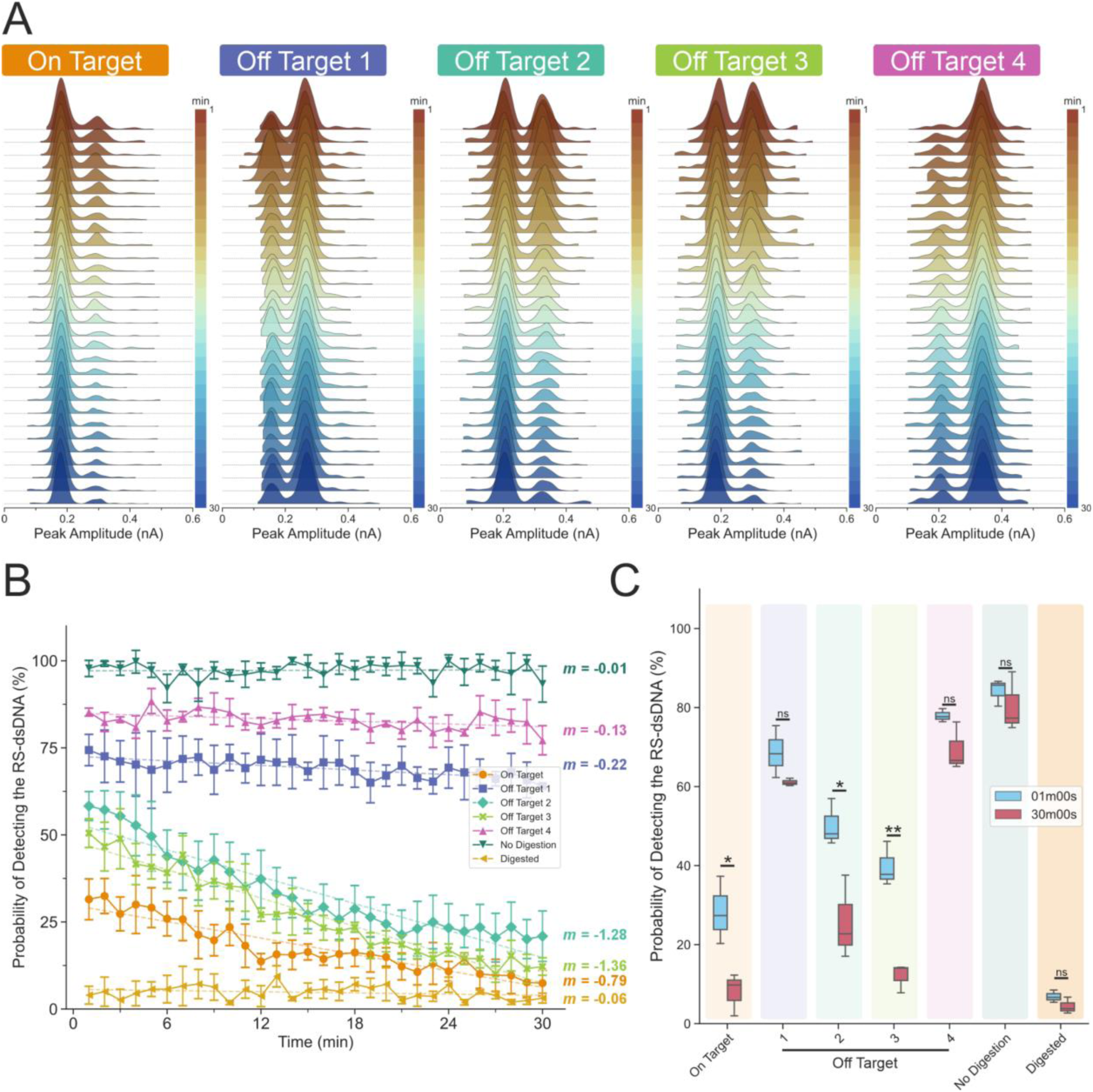
Measuring the activity of the Cas9 endonuclease with nanopore. **(A),** Ridgeline plot showing the KDE curve of the translocation experiment at each min for the 5 tested on target and off target crRNA sequence. **(B),** Probability of detecting 3kbp dsDNA as a function of the digestion time for all the variants tested and controls. *m* is the slope after fitting with the linear fit to the experimental data. Error bars are standard error of the mean. **(C),** Box plot comparing the probability of detecting the RS-dsDNA at 1 min and at 30 min. Two-tailed unpaired T test was used to test the differences between the distribution of the probabilities to detect RS-dsDNA at 1 min and at 30 min, there are significant differences for the on target, off target 2 and off target 3 at 1 min and at 30 min. The calculated values for on target at 1 min is 28.31±8.55% and 8.01±5.37% at 30 min respectively, for off target 1 at 1 min is 68.67±6.57% and 61.08±0.94% respectively, for off target 2 at 1 min is 50.25±5.93% and 25.79±10.6% respectively, for off target 3 at 1 min is 39.76±5.65% and 12.04±3.67% respectively, for off target 4 at 1 min is 77.95±1.67% and 69.36±6.11% respectively, for no digestion at 1 min is 84.22±3.38% and 80.44±7.56% respectively, for digested at 1 min is 8.52±1.72% and 4.39±2.07% respectively. (ns, not significant; *, P ˂0.05; **, P˂0.005; data assume normal distribution; Levene’s test (P>0.05) indicates data are homoscedasticity; N=3, two tailed T-test).

We quantified the probabilities of detecting the RS-dsDNA using all variants of crRNA-Cas9 RNP mixtures as well as the no digestion (S.Figure 14) and digested (S.Figure 15) (pre-incubated with on target Cas9 RNP for 30 min prior measurement) controls (Figure 6B). The no digestion and Cas9 RNP digested controls showed that the probabilities to detect 3.0 kbp RS-dsDNA were at near 90% and at near 10%, respectively, throughout the experiment. The linear fitted slope had a minimum value of *m*>-0.1 over the measurement time and there was no significant difference found between these controls at 1 min and the 30 min (Figure 6C). The off-target 1 and 4 variants showed a slow downward trend in the probability of detecting RS-dsDNA, with slope changes of *m*<-0.1 (Figure 6B). The were no significant differences in the probabilities of detecting RS-dsDNA between the 1 min and 30 min for these crRNA variants (Figure 6C), indicating a near negligible progression of the cleavage.

For the off target 2 and 3 variants, the slopes were fitted to be *m*<-1, in sharp contrast to other variants and indicated faster cleavage activities (Figure 6B). Additionally, there were significant differences in the probabilities at 1 min and 30 min where both variants showed lower probabilities of detecting the RS-dsDNA (Figure 6C). Overall, the two variants were more catalytically active and cleaved at similar speed, off-target 2 variant slowed the reaction rate significantly more than off-target 3 (S.Figure 16), despite the mutations were essentially the same sequence swap from rA-dT to rG-dT, and our study revealed that the position upstream of PAM played a significant role in determining the cleavage kinetics. Lastly, the on-target Cas9 RNPs cleaved the RS-dsDNA quite quickly such that there was only about a 25% probability of detecting the uncleaved RS-dsDNA by the time the experiment started. It then cleaved the remaining population at a slower rate with a slope of −0.8 through the remainder of the experiment.

Mismatches in the seed or PAM proximal region typically interrupt R-loop formation and often lead to higher dissociation rates of the Cas9 RNP from the target DNA, reducing or eliminating cleavage activity as well ^26,64,67^. Additionally, the cleavage activities could be lower due to distortions introducing steric hindrance between the HNH endonuclease domain and the heteroduplex. The variants we introduced form non form noncanonical base pairing including wobble base pairing (rU-dG, rG-dT; position 1-3) ^74^ and non-isosteric pyrimidine-pyrimidine base pairing (rU-dC, position 4) ^75^ when constrained within the Cas9 RNPs complexes ^64^. In our experiments, mismatches in the seed region at position 1 and 4 lead to suppressed cleavage, However, we have observed off-target or mismatch tolerance at positions 2 and 3 that result in cleavage kinetics that are slightly slower but comparable to the on target Cas9 RNP. Non canonical binding (wobble base pairing) may contribute to stabilizing the binding in these cases. At position 1 there is also a potential wobble base pairing (rU-dG), however, due to the position of the mismatch, the sequence rigidity is expected to hinder the binding of the wobble base pair which could contribute to the suppression of the cleavage activity due to steric hindrance ^26,64,67^. These measurements demonstrate the potential of our nanopore sensor to rapidly provide detailed kinetic information on the activity of enzymes in physiological conditions.

### Conclusion

In this study, we developed and validated a solid-state nanopore based kinetic sensing system to monitor the activities of endonucleases such as the restriction enzyme, SwaI, and Cas9 RNPs in physiologically relevant salt conditions. Our nanopore sensor effectively distinguishes between the reactant (RS-dsDNA) and cleavage products (1.5 kbp dsDNA) due to the molecular weight differences and thus is able to quantify the population with single molecule resolution. As the digestion progresses, the population differences were monitored in real-time, and an observable depletion of the RS-dsDNA could be seen in both the restriction enzyme and Cas9 RNP cases. To facilitate analysis, the probability density functions of the populations were estimated through non-parametric kernel density estimation, and estimated the population as probabilities for comparison between different conditions and variants.

We have presented a proof of concept of a single molecule sensing system that enables real-time label-free measurements of enzyme activity. Several improvements to the measurement and analysis system would allow us to further expand our applications. For example, the hardware could be improved by implementing a temperature controlling unit, to allow us to monitor temperature sensitive endonucleases. This improvement could also allow us to dynamically control the temperature of the electrolyte bath to either enhance, reduce, or abolish the endonuclease activity. Secondly, it is hard to perform liquid exchange or mixing after backfilling the current glass capillaries. To this end, wider outer diameter quartz capillaries tube could be used to fabricate the nanopores to facilitate solution mixing in the future. Thirdly, the boundary selection for the quantification of the probability was fixed and applied across all the PDFs. However, to allow for a longer experiment, the boundary selection and calculation for the data analysis could dynamically adapt to the baseline current level.

Unlike most solid-state nanopore approaches, the method developed here takes advantage of the unique properties of the polymer electrolyte bath measurement system to eliminate the need for extremely high concentrations of salt to improve the detection resolution ^31,34^, thus allowing us to monitor the activities of restriction enzyme and Cas9 RNP endonucleases at physiological relevant salt conditions. Since the analyte signal does not depend on the properties of the buffer, as we previously demonstrated ^31^, salt sensitive analytes such as intrinsically disordered proteins ^76,77^ can be analysed and monitored over time at single molecule resolution with the nanopore. For example, the aggregation of amyloid materials ^35,78^ under different salt conditions can be monitored as aggregation propensity is different under different ionic strength ^79–84^. While the method developed here focuses on monitoring the transition of reactant to product, the same method can, in principle, be applied to study other reactions such as full digestion of materials (such as full or partial digestion of nucleic acid by DNase) ^85^ and the emergence of larger biomolecules during overtime (such as oligomer aggregation) ^79–84^.

## Methods

### Solid-state Nanopore fabrication and measurement

The glass solid-state nanopore (nanopipette) was fabricated by the SU-P2000 laser puller (World Precision Instruments). Quartz capillaries of 1.0 mm outer diameter and 0.5 mm inner diameter with filament (QF100-50-7.5; Sutter Instrument) were used for the nanopore fabrication. A two-line protocol was used: line 1, HEAT 750/FIL 4/VEL 30/DEL 150/PUL 80, followed by line 2, HEAT 725/FIL 3/VEL 40/DEL 135/PUL 180. The pulling protocol is instrument specific, and there is variation between pullers. The nanopore dimension was confirmed by scanning electron microscopy.

The standard translocation experiment follows a similar procedure from the previous publication. The analyte filled nanopore was fitted with a Ag/AgCl working electrode and immersed into the polymer electrolyte bath with a Ag/AgCl reference electrode. Ionic current trace was recorded using the MultiClamp 700B patch-clamp amplifier (Molecular Devices) in voltage-clamp mode. The sampling bandwidth was approximately 52 kHz. The signal was filtered using a low-pass filter at 20 kHz setting, digitized with a Digidata 1550B (Molecular Devices) at a 100 kHz (10 μs) sampling rate. The software used for recording was pClamp 10 (Molecular Devices). For translocation events analysis, the threshold level was defined at least 10 sigma away from the baseline, only events that were above this threshold would be identified as the translocation of the molecule. The analysis script can be accessed here: https://github.com/chalmers4c/Nanopore_event_detection.

### Polymer electrolyte bath generation

The generation of the PEG polymer electrolyte bath follows the exact same procedure as previously published study. Briefly, to generate 10 ml of the 50% (w/v) PEG 35K KCl electrolyte bath, 5 g of PEG 35K (94646; Sigma-Aldrich) was mixed with 1 ml of 1 M KCl (A11662.0B; Thermo Fisher) and 4 ml of ddH_2_O. The mixture was then incubated inside an 85 °C oven for 2 hours followed by overnight incubation at 37 °C. All the electrolyte baths were discarded one month after the generation.

### Kinetic Translocation experiment

Prior to the measurement, the RE-dsDNA, the restriction enzyme SwaI with the restriction digestion buffer was mixed so that the RS-dsDNA, SwaI and the buffer was at 10 nM, 5 units and 1× respectively. The mixture was immediately loaded into the glass nanopore and immersed into the polymer electrolyte bath, with a Ag/AgCl working electrode fitted into the nanopore.

From mixing the reactants to start of measurement, we estimated a delay of 1 min. We applied a waveform composed of 3.5 seconds of +100mV followed by 6 seconds of −700mV. The single molecule events recorded between 4 to 9 seconds of each trace were used for all analysis.

Supporting information contains more details on the method for the analysis of the trace. A custom written python script was used for the calculation of the boundaries and the AUC. The fitting was carried out with python, the linear fit was calculated by sum of least squares method.

### CRISPR-Cas9 ribonucleoproteins complex assembly and reaction

The recombinant S. pyogenes Cas9 was used (1081058; IDTDNA) throughout the study. The tracrRNA (1072532; IDTDNA) and crRNA were synthesized and provided by IDTDNA. The tracrRNA and crRNA was mixed and diluted with the duplex buffer (30 mM HEPES, pH 7.5; 100 mM potassium acetate; 11-01-03-01; IDTDNA), the final mixture contained 40 µM of tracrRNA and 40 µM of crRNA. The mixture was incubated at 95°C for 5 min and then at 25°C until use. To assemble the Cas9 RNPs, the RNA mixture after the incubation was mixed with the Cas9 proteins and diluted with digestion buffer (111 mM HEPES at pH 8.0, 6 mM MgCl_2_, 111 mM NaCl), so that the final mixture contained 40 µM tracrRNA, 40 µM of crRNA and 18.6 µM of Cas9 proteins, followed by incubation at 25°C for 30 min, and store at 4°C until use. For longer storage, the Cas9 RNPs were snap frozen with liquid nitrogen and stored at −80°C. For kinetic translocation experiment, the Cas9 RNPs were mixed with RS-dsDNA to a final concentration of 1000 nM Cas9 RNPs and 10 nM dsDNA (100:1 Cas9 RNP to dsDNA ratio) and loaded into the nanopore immediately prior to use. The measurement set-ups and routines were the same as kinetic translocation experiment session.

## Data availability

All the translocation trace data supporting this work can be freely accessed via the University of Leeds data repository: https://doi.org/10.5518/1454

## Supporting Information

Supporting information include detailed information: supporting note on polymer electrolyte modified solid-state nanopore mechanism, generation of the RS-dsDNA and the sequence data, the generation of the RS-dsDNA, restriction digestion of the RS-dsDNA, Cas9 related crRNA sequence and digestion, kernel density estimation and probability calculations, and relevant supporting figures.

## Author Contributions

C.C. designed and performed all experiments. C.C. analysed all data. C.C. illustrated all schematics. N.E.W., E.E.T. and P.A. provided the concept of Cas9 detection and helped data analysis. P.A. helped with data analysis and acquired the funding. All authors wrote and corrected the manuscript.

## Acknowledgement

We thank the group of bioelectronics member and Professor Christoph Wälti of University of Leeds for providing insightful feedback. We thank Dr Alexander Kulak of University of Leeds for imaging the nanopore with the scanning electron microscopy. C.C. thanks the CP 1919 pulsar and Joy Division’s “Unknown Pleasures” album cover for the inspiration that directly led to the creation of the ridgeline plots for the visualisation of the enzyme digestion kinetics.

## Abbreviation

AUC: Area under the curve

KDE: kernel density estimation

PEG: poly(ethylene) glycol

PCR: polymerase chain reaction

RS-dsDNA: Restriction site containing dsDNA

CRISPR: clustered regularly interspaced short palindromic repeats

PAM: Protospacer adjacent motif

RNP: Ribonucleoprotein

