## Supporting Information for "Solid-State Nanopore Real Time Assay for Monitoring Cas9 Endonuclease Reactivity"

### Note 1: Polymer electrolyte modified Solid-state nanopore mechanism

In contrast to most solid-state nanopore systems where the dwell time component is responsible for the separation on the length of dsDNA and the current amplitude component is resistive instead of conductive [1-3], these differences are due to the polymer electrolyte. Based on the conductive current amplitude alone, we previously reported that different sizes of the dsDNA ranging from 0.7 to 7 kbp could be reliably detected and distinguished [4]. Under this modified polymer electrolyte condition, the nanopore becomes permselective to cations, as evidenced by the extreme ion current rectification in the voltammograms (S.Figure 2, 6). This enhances the sensitivity of the nanopore towards dsDNA which is known to have a highly charged ionic cloud [5-7], and leads to conductive translocation events[8,9]. The use of a polymer electrolyte in the trans chamber and the standard electrolyte solution inside the nanopore creates a physical interface at the nanopore. The translocation of dsDNA from cis to trans chambers cause a mechanical disruption of this interface, leading to an accumulation of ions at the aperture which corresponds to a temporal enhanced conductivity in the system. Importantly, the length of the interface protrusion is linearly correlated with the length of the dsDNA translocating from cis to trans chamber [4].

### Method

Unless otherwise specified, all chemicals were obtained from Sigma-Aldrich at ACS grade.

#### Generation of the RS-dsDNA

The generation of the Restriction site containing dsDNA (RS-dsDNA) involves multiple steps.

##### Fragment 1 and Fragment 2 generation

The m13mp18 ssDNA (N4040S; NEB) was used as the PCR template to generate two separate fragments: Fragment 1 and Fragment 2. The primers used for the PCR reaction contains the newly introduced SwaI restriction site region. The LongAmp® *Taq* PCR kit (E5200S; NEB) was used to produce both the Fragment 1 and Fragment 2. The PCR reaction mixtures were composed of 1 ng m13mp18 ssDNA, 1X LongAmp® *Taq* Reaction buffer, 300 μM dNTPs, 0.4 μM forward primer, 0.4 μM reverse primer and 1 units/10 μl of LongAmp® *Taq* DNA polymerase, diluted in nuclease-free ddH$_2$O. The PCR reaction mixtures were subjected to the following PCR cycles in the Mastercycler Nexus X2 (Eppendorf): 94°C for 4 mins, 30 cycles of 94°C for 30 seconds, followed by 56°C for 30 seconds and then 65°C for 2 mins and 20 seconds, after the 30 cycles, the mixtures were held at 65°C for 10 mins, then dropped down to 25°C for infinite hold. After the PCR reaction, the mixtures were cleaned up with the Monarch® PCR & DNA Cleanup Kit (T1030; NEB). The concentration and purity of the PCR products were checked with absorption spectroscopy at $A_{260}$.

##### Assembled the Fragment 1 and Fragment 2

The Fragment 1 and Fragment were joined by Gibson assembly method [10]. The NEBuilder® HiFi DNA Assembly (E2621; NEB) was used. In a 20 μl reaction mixture, 10 ng of Fragment 1 and 10 ng of Fragment 2 were mixed with 10 μl of the NEBuilder HiFi DNA Assembly Master Mix, and diluted with nuclease-free ddH$_2$O. The reaction mixture was incubated inside the Mastercycler Nexus X2 at 50°C for 60 mins, and then dropped to 4°C for infinite hold. The reaction mixture was cleaned and purified with the Monarch® PCR & DNA Cleanup Kit, followed by analysis with absorption spectroscopy at $A_{260}$. PCR was used to amplify the quantity of the assembled products. The LongAmp® *Taq* PCR kit was used with the following components in the final PCR reaction mixture: 5 ng of assembled products, 1X LongAmp® *Taq* Reaction buffer, 300 μM dNTPs, 0.4 μM forward primer, 0.4 μM reverse primer and 1 units/10 μl of LongAmp® *Taq* DNA polymerase, diluted in nuclease-free ddH$_2$O. The PCR protocol was as follows: 94°C for 4 mins, 30 cycles of 94°C for 30 seconds, followed by 56°C for 30 seconds and then 65°C for 2 mins and 20 seconds, after the 30 cycles, the mixtures were held at 65°C for 10 mins, then dropped down to 25°C for infinite hold inside the Mastercycler Nexus X2. The PCR products were purified with the Monarch® PCR & DNA Cleanup Kit and analysed with absorption spectroscopy at $A_{260}$.

The purified products were analysed with agarose gel electrophoresis. The fragments corresponding to the assembled products were cut with a blade, isolated from the rest of the fragments and purified with the Monarch® DNA Gel Extraction Kit (T1020). The purified assembled product was checked with absorption spectroscopy at $A_{260}$.

##### RS-dsDNA production

The 3 kbp RS-dsDNA was produced by PCR using the isolated purified assembled product as the template. The KAPA HiFi HotStart PCR Kit (KR0369; Roche) was used, the reaction component contained 1 ng of the purified assembled product, 1X KAPA HiFi HotStart buffer, 300 µM dNTPs, 0.3 µM forward primer, 0.3 reverse primer, 1 unit/10 µl KAPA HiFi HotStart PCR polymerase and diluted in nuclease-free $ddH_2O$. The Mastercycler Nexus X2 was used to perform the PCR reaction with the following parameters: 95°C for 3 mins, 26 cycles of 98°C for 20 seconds, followed by 62°C for 15 seconds and then 72°C for 4 mins, after the 26 cycles, the mixtures were held at 72°C for 4 mins, then dropped down to 25°C for infinite hold. The PCR products were purified via PEG precipitation. To increase the quantity of the final products, multiple of the same PCR reactions were pooled into a single tube. The PEG precipitation buffer was composed of 2.5M NaCl and 20% (w/v) PEG 8000 (89510; Sigma-Aldrich), the precipitation buffer was mixed with the PCR products at 1:1 volume to volume ratio, followed by a brief vortex and incubate at 37°C for 30 mins. The mixture was then centrifuged for 15 mins at 20,000 rpm. The supernatant was removed by pipette and the pellet was resuspended and washed with 150 µl of 80% (v/v) ice cold ethanol, followed by centrifugation at 20,000 rpm for 10 mins. The supernatant was discarded and the pellet was washed again with 96% (v/v) ice cold ethanol, followed by centrifugation again at 20,000 rpm for 10 mins. The supernatant was discarded and left to dry on bench for 10 mins. The pellet was resuspended in 20 µl of 1X TE buffer (AM9858; Thermo Scientific) and checked with absorption spectroscopy at $A_{260}$.

#### Sequence

##### Fragment 1 and Fragment 2 related

The sequence for the m13mp18:

5′–

AATGCTACTACTATTAGTAGAATTGATGCCACCTTTTCAGCTCGCGCCCCAAATGAAAATATAGCTAA
ACAGGTTATTGACCATTTGCGAAATGTATCTAATGGTCAAACTAAATCTACTCGTTCGCAGAATTGGG
AATCAACTGTTATATGGAATGAAACTTCCAGACACCGTACTTTAGTTGCATATTTAAAACATGTTGAG

CTACAGCATTATATTCAGCAATTAAGCTCTAAGCCATCCGCAAAAATGACCTCTTATCAAAAGGAGCA
ATTAAAGGTACTCTCTAATCCTGACCTGTTGGAGTTTGCTTCCGGTCTGGTTCGCTTTGAAGCTCGAA
TTAAAACGCGATATTTGAAGTCTTTCGGGCTTCCTCTTAATCTTTTTGATGCAATCCGCTTTGCTTCT
GACTATAATAGTCAGGGTAAAGACCTGATTTTTGATTTATGGTCATTCTCGTTTTCTGAACTGTTTAA
AGCATTTGAGGGGGATTCAATGAATATTTATGACGATTCCGCAGTATTGGACGCTATCCAGTCTAAAC
ATTTTACTATTACCCCCTCTGGCAAAACTTCTTTTGCAAAAGCCTCTCGCTATTTTGGTTTTTATCGT
CGTCTGGTAAACGAGGGTTATGATAGTGTTGCTCTTACTATGCCTCGTAATTCCTTTTGGCGTTATGT
ATCTGCATTAGTTGAATGTGGTATTCCTAAATCTCAACTGATGAATCTTTCTACCTGTAATAATGTTG
TTCCGTTAGTTCGTTTTATTAACGTAGATTTTTCTTCCCAACGTCCTGACTGGTATAATGAGCCAGTT
CTTAAAATCGCATAAGGTAATTCACAATGATTAAAGTTGAAATTAAACCATCTCAAGCCCAATTTACT
ACTCGTTCTGGTGTTTCTCGTCAGGGCAAGCCTTATTCACTGAATGAGCAGCTTTGTTACGTTGATTT
GGGTAATGAATATCCGGTTCTTGTCAAGATTACTCTTGATGAAGGTCAGCCAGCCTATGCGCCTGGTC
TGTACACCGTTCATCTGTCCTCTTTCAAAGTTGGTCAGTTCGGTTCCCTTATGATTGACCGTCTGCGC
CTCGTTCCGGCTAAGTAACATGGAGCAGGTCGCGGATTTCGACACAATTTATCAGGCGATGATACAAA
TCTCCGTTGTACTTTGTTTCGCGCTTGGTATAATCGCTGGGGGTCAAAGATGAGTGTTTTAGTGTATT
CTTTTGCCTCTTTCGTTTTAGGTTGGTGCCTTCGTAGTGGCATTACGTATTTTACCCGTTTAATGGAA
ACTTCCTCATGAAAAAGTCTTTAGTCCTCAAAGCCTCTGTAGCCGTTGCTACCCTCGTTCCGATGCTG
TCTTTCGCTGCTGAGGGTGACGATCCCGCAAAAGCGGCCTTTAACTCCCTGCAAGCCTCAGCGACCGA
ATATATCGGTTATGCGTGGGCGATGGTTGTTGTCATTGTCGGCGCAACTATCGGTATCAAGCTGTTTA
AGAAATTCACCTCGAAAGCAAGCTGATAAACCGATACAATTAAAGGCTCCTTTTGGAGCCTTTTTTTT
GGAGATTTTCAACGTGAAAAAATTATTATTCGCAATTCCTTTAGTTGTTCCTTTCTATTCTCACTCCG
CTGAAACTGTTGAAAGTTGTTTAGCAAAATCCCATACAGAAAATTCATTTACTAACGTCTGGAAAGAC
GACAAAACTTTAGATCGTTACGCTAACTATGAGGGCTGTCTGTGGAATGCTACAGGCGTTGTAGTTTG
TACTGGTGACGAAACTCAGTGTTACGGTACATGGGTTCCTATTGGGCTTGCTATCCCTGAAAATGAGG
GTGGTGGCTCTGAGGGTGGCGGTTCTGAGGGTGGCGGTTCTGAGGGTGGCGGTACTAAACCTCCTGAG
TACGGTGATACACCTATTCCGGGCTATACTTATATCAACCCTCTCGACGGCACTTATCCGCCTGGTAC
TGAGCAAAACCCCGCTAATCCTAATCCTTCTCTTGAGGAGTCTCAGCCTCTTAATACTTTCATGTTTC
AGAATAATAGGTTCCGAAATAGGCAGGGGGCATTAACTGTTTATACGGGCACTGTTACTCAAGGCACT
GACCCCGTTAAAACTTATTACCAGTACACTCCTGTATCATCAAAAGCCATGTATGACGCTTACTGGAA
CGGTAAATTCAGAGACTGCGCTTTCCATTCTGGCTTTAATGAGGATTTATTTGTTTGTGAATATCAAG
GCCAATCGTCTGACCTGCCTCAACCTCCTGTCAATGCTGGCGGCGGCTCTGGTGGTGGTTCTGGTGGC
GGCTCTGAGGGTGGTGGCTCTGAGGGTGGCGGTTCTGAGGGTGGCGGCTCTGAGGGAGGCGGTTCCGG
TGGTGGCTCTGGTTCCGGTGATTTTGATTATGAAAAGATGGCAAACGCTAATAAGGGGGCTATGACCG
AAAATGCCGATGAAAACGCGCTACAGTCTGACGCTAAAGGCAAACTTGATTCTGTCGCTACTGATTAC
GGTGCTGCTATCGATGGTTTCATTGGTGACGTTTCCGGCCTTGCTAATGGTAATGGTGCTACTGGTGA
TTTTGCTGGCTCTAATTCCCAAATGGCTCAAGTCGGTGACGGTGATAATTCACCTTTAATGAATAATT
TCCGTCAATATTTACCTTCCCTCCCTCAATCGGTTGAATGTCGCCCTTTTGTCTTTGGCGCTGGTAAA

CCATATGAATTTTCTATTGATTGTGACAAAATAAACTTATTCCGTGGTGTCTTTGCGTTTCTTTTATA
TGTTGCCACCTTTATGTATGTATTTTCTACGTTTGCTAACATACTGCGTAATAAGGAGTCTTAATCAT
GCCAGTTCTTTTGGGTATTCCGTTATTATTGCGTTTCCTCGGTTTCCTTCTGGTAACTTTGTTCGGCT
ATCTGCTTACTTTTCTTAAAAAGGGCTTCGGTAAGATAGCTATTGCTATTTCATTGTTTCTTGCTCTT
ATTATTGGGCTTAACTCAATTCTTGTGGGTTATCTCTCTGATATTAGCGCTCAATTACCCTCTGACTT
TGTTCAGGGTGTTCAGTTAATTCTCCCGTCTAATGCGCTTCCCTGTTTTTATGTTATTCTCTCTGTAA
AGGCTGCTATTTTCATTTTTGACGTTAAACAAAAAATCGTTTCTTATTTGGATTGGGATAAATAATAT
GGCTGTTTATTTTGTAACTGGCAAATTAGGCTCTGGAAAGACGCTCGTTAGCGTTGGTAAGATTCAGG
ATAAAATTGTAGCTGGGTGCAAAATAGCAACTAATCTTGATTTAAGGCTTCAAAACCTCCCGCAAGTC
GGGAGGTTCGCTAAAACGCCTCGCGTTCTTAGAATACCGGATAAGCCTTCTATATCTGATTTGCTTGC
TATTGGGCGCGGTAATGATTCCTACGATGAAAATAAAAACGGCTTGCTTGTTCTCGATGAGTGCGGTA
CTTGGTTTAATACCCGTTCTTGGAATGATAAGGAAAGACAGCCGATTATTGATTGGTTTCTACATGCT
CGTAAATTAGGATGGGATATTATTTTTCTTGTTCAGGACTTATCTATTGTTGATAAACAGGCGCGTTC
TGCATTAGCTGAACATGTTGTTTATTGTCGTCGTCTGGACAGAATTACTTTACCTTTTGTCGGTACTT
TATATTCTCTTATTACTGGCTCGAAAATGCCTCTGCCTAAATTACATGTTGGCGTTGTTAAATATGGC
GATTCTCAATTAAGCCCTACTGTTGAGCGTTGGCTTTATACTGGTAAGAATTTGTATAACGCATATGA
TACTAAACAGGCTTTTTCTAGTAATTATGATTCCGGTGTTTATTCTTATTTAACGCCTTATTTATCAC
ACGGTCGGTATTTCAAACCATTAAATTTAGGTCAGAAGATGAAATTAACTAAAATATATTTGAAAAAG
TTTTCTCGCGTTCTTTGTCTTGCGATTGGATTTGCATCAGCATTTACATATAGTTATATAACCCAACC
TAAGCCGGAGGTTAAAAAGGTAGTCTCTCAGACCTATGATTTTGATAAATTCACTATTGACTCTTCTC
AGCGTCTTAATCTAAGCTATCGCTATGTTTTCAAGGATTCTAAGGGAAAATTAATTAATAGCGACGAT
TTACAGAAGCAAGGTTATTCACTCACATATATTGATTTATGTACTGTTTCCATTAAAAAAGGTAATTC
AAATGAAATTGTTAAATGTAATTAATTTTGTTTTCTTGATGTTTGTTTCATCATCTTCTTTTGCTCAG
GTAATTGAAATGAATAATTCGCCTCTGCGCGATTTTGTAACTTGGTATTCAAAGCAATCAGGCGAATC
CGTTATTGTTTCTCCCGATGTAAAAGGTACTGTTACTGTATATTCATCTGACGTTAAACCTGAAAATC
TACGCAATTTCTTTATTTCTGTTTTACGTGCAAATAATTTTGATATGGTAGGTTCTAACCCTTCCATT
ATTCAGAAGTATAATCCAAACAATCAGGATTATATTGATGAATTGCCATCATCTGATAATCAGGAATA
TGATGATAATTCCGCTCCTTCTGGTGGTTTCTTTGTTCCGCAAAATGATAATGTTACTCAAACTTTTA
AAATTAATAACGTTCGGGCAAAGGATTTAATACGAGTTGTCGAATTGTTTGTAAAGTCTAATACTTCT
AAATCCTCAAATGTATTATCTATTGACGGCTCTAATCTATTAGTTGTTAGTGCTCCTAAAGATATTTT
AGATAACCTTCCTCAATTCCTTTCAACTGTTGATTTGCCAACTGACCAGATATTGATTGAGGGTTTGA
TATTTGAGGTTCAGCAAGGTGATGCTTTAGATTTTTCATTTGCTGCTGGCTCTCAGCGTGGCACTGTT
GCAGGCGGTGTTAATACTGACCGCCTCACCTCTGTTTTATCTTCTGCTGGTGGTTCGTTCGGTATTTT
TAATGGCGATGTTTTAGGGCTATCAGTTCGCGCATTAAAGACTAATAGCCATTCAAAAATATTGTCTG
TGCCACGTATTCTTACGCTTTCAGGTCAGAAGGGTTCTATCTCTGTTGGCCAGAATGTCCCTTTTATT
ACTGGTCGTGTGACTGGTGAATCTGCCAATGTAAATAATCCATTTCAGACGATTGAGCGTCAAAATGT
AGGTATTTCCATGAGCGTTTTTCCTGTTGCAATGGCTGGCGGTAATATTGTTCTGGATATTACCAGCA

AGGCCGATAGTTTGAGTTCTTCTACTCAGGCAAGTGATGTTATTACTAATCAAAGAAGTATTGCTACA
ACGGTTAATTTGCGTGATGGACAGACTCTTTTACTCGGTGGCCTCACTGATTATAAAAACACTTCTCA
GGATTCTGGCGTACCGTTCCTGTCTAAAATCCCTTTAATCGGCCTCCTGTTTAGCTCCCGCTCTGATT
CTAACGAGGAAAGCACGTTATACGTGCTCGTCAAAGCAACCATAGTACGCGCCCTGTAGCGGCGCATT
AAGCGCGGCGGGTGTGGTGGTTACGCGCAGCGTGACCGCTACACTTGCCAGCGCCCTAGCGCCCGCTC
CTTTCGCTTTCTTCCCTTCCTTTCTCGCCACGTTCGCCGGCTTTCCCCGTCAAGCTCTAAATCGGGGG
CTCCCTTTAGGGTTCCGATTTAGTGCTTTACGGCACCTCGACCCCAAAAAACTTGATTTGGGTGATGG
TTCACGTAGTGGGCCATCGCCCTGATAGACGGTTTTTCGCCCTTTGACGTTGGAGTCCACGTTCTTTA
ATAGTGGACTCTTGTTCCAAACTGGAACAACACTCAACCCTATCTCGGGCTATTCTTTTGATTTATAA
GGGATTTTGCCGATTTCGGAACCACCATCAAACAGGATTTTCGCCTGCTGGGGCAAACCAGCGTGGAC
CGCTTGCTGCAACTCTCTCAGGGCCAGGCGGTGAAGGGCAATCAGCTGTTGCCCGTCTCACTGGTGAA
AAGAAAAACCACCCTGGCGCCCAATACGCAAACCGCCTCTCCCCGCGCGTTGGCCGATTCATTAATGC
AGCTGGCACGACAGGTTTCCCGACTGGAAAGCGGGCAGTGAGCGCAACGCAATTAATGTGAGTTAGCT
CACTCATTAGGCACCCCAGGCTTTACACTTTATGCTTCCGGCTCGTATGTTGTGTGGAATTGTGAGCG
GATAACAATTTCACACAGGAAACAGCTATGACCATGATTACGAATTCGAGCTCGGTACCCGGGGATCC
TCTAGAGTCGACCTGCAGGCATGCAAGCTTGGCACTGGCCGTCGTTTTACAACGTCGTGACTGGGAAA
ACCCTGGCGTTACCCAACTTAATCGCCTTGCAGCACATCCCCCTTTCGCCAGCTGGCGTAATAGCGAA
GAGGCCCGCACCGATCGCCCTTCCCAACAGTTGCGCAGCCTGAATGGCGAATGGCGCTTTGCCTGGTT
TCCGGCACCAGAAGCGGTGCCGGAAAGCTGGCTGGAGTGCGATCTTCCTGAGGCCGATACTGTCGTCG
TCCCCTCAAACTGGCAGATGCACGGTTACGATGCGCCCATCTACACCAACGTGACCTATCCCATTACG
GTCAATCCGCCGTTTGTTCCCACGGAGAATCCGACGGGTTGTTACTCGCTCACATTTAATGTTGATGA
AAGCTGGCTACAGGAAGGCCAGACGCGAATTATTTTTGATGGCGTTCCTATTGGTTAAAAAATGAGCT
GATTTAACAAAAATTTAATGCGAATTTTAACAAAATATTAACGTTTACAATTTAAATATTTGCTTATA
CAATCTTCCTGTTTTTGGGGCTTTTCTGATTATCAACCGGGGTACATATGATTGACATGCTAGTTTTA
CGATTACCGTTCATCGATTCTCTTGTTTGCTCCAGACTCTCAGGCAATGACCTGATAGCCTTTGTAGA
TCTCTCAAAAATAGCTACCCTCTCCGGCATTAATTTATCAGCTAGAACGGTTGAATATCATATTGATG
GTGATTTGACTGTCTCCGGCCTTTCTCACCCTTTTGAATCTTTACCTACACATTACTCAGGCATTGCA
TTTAAAATATATGAGGGTTCTAAAAATTTTTATCCTTGCGTTGAAATAAAGGCTTCTCCCGCAAAAGT
ATTACAGGGTCATAATGTTTTTGGTACAACCGATTTAGCTTTATGCTCTGAGGCTTTATTGCTTAATT
TTGCTAATTCTTTGCCTTGCCTGTATGATTTATTGGATGTT

-3′

The forward primer for Fragment 1:

5′-**ATTGGGATTT**AAATCGATGAGTGCGGTAC-3′

The reverse primer for Fragment 1:

5′-GCTTTGACGAGCACGTATAAC-3′

The forward primer for Fragment 2:

5′-CGCAAAAGCGGCCTTTAAC-3′

The reverse primer for Fragment 2:

5′-**TCATCGATTT** AAATCCCAATAGCAAGCAAATC-3′

#### Assembled Fragment 1 and Fragment 2 related

The assembled product sequence:

5′-

CGCAAAAGCGGCCTTTAACTCCCTGCAAGCCTCAGCGACCGAATATATCGGTTATGCGTGGGCGATGG
TTGTTGTCATTGTCGGCGCAACTATCGGTATCAAGCTGTTTAAGAAATTCACCTCGAAAGCAAGCTGA
TAAACCGATACAATTAAAGGCTCCTTTTGGAGCCTTTTTTTTGGAGATTTTCAACGTGAAAAAATTAT
TATTCGCAATTCCTTTAGTTGTTCCTTTCTATTCTCACTCCGCTGAAACTGTTGAAAGTTGTTTAGCA
AAATCCCATACAGAAAATTCATTTACTAACGTCTGGAAAGACGACAAAACTTTAGATCGTTACGCTAA
CTATGAGGGCTGTCTGTGGAATGCTACAGGCGTTGTAGTTTGTACTGGTGACGAAACTCAGTGTTACG
GTACATGGGTTCCTATTGGGCTTGCTATCCCTGAAAATGAGGGTGGTGGCTCTGAGGGTGGCGGTTCT
GAGGGTGGCGGTTCTGAGGGTGGCGGTACTAAACCTCCTGAGTACGGTGATACACCTATTCCGGGCTA
TACTTATATCAACCCTCTCGACGGCACTTATCCGCCTGGTACTGAGCAAAACCCCGCTAATCCTAATC
CTTCTCTTGAGGAGTCTCAGCCTCTTAATACTTTCATGTTTCAGAATAATAGGTTCCGAAATAGGCAG
GGGGCATTAACTGTTTATACGGGCACTGTTACTCAAGGCACTGACCCCGTTAAAACTTATTACCAGTA
CACTCCTGTATCATCAAAAGCCATGTATGACGCTTACTGGAACGGTAAATTCAGAGACTGCGCTTTCC
ATTCTGGCTTTAATGAGGATTTATTTGTTTGTGAATATCAAGGCCAATCGTCTGACCTGCCTCAACCT
CCTGTCAATGCTGGCGGCGGCTCTGGTGGTGGTTCTGGTGGCGGCTCTGAGGGTGGTGGCTCTGAGGG
TGGCGGTTCTGAGGGTGGCGGCTCTGAGGGAGGCGGTTCCGGTGGTGGCTCTGGTTCCGGTGATTTTG
ATTATGAAAAGATGGCAAACGCTAATAAGGGGGCTATGACCGAAAATGCCGATGAAAACGCGCTACAG
TCTGACGCTAAAGGCAAACTTGATTCTGTCGCTACTGATTACGGTGCTGCTATCGATGGTTTCATTGG
TGACGTTTCCGGCCTTGCTAATGGTAATGGTGCTACTGGTGATTTTGCTGGCTCTAATTCCCAAATGG
CTCAAGTCGGTGACGGTGATAATTCACCTTTAATGAATAATTTCCGTCAATATTTACCTTCCCTCCCT
CAATCGGTTGAATGTCGCCCTTTTGTCTTTGGCGCTGGTAAACCATATGAATTTTCTATTGATTGTGA
CAAAATAAACTTATTCCGTGGTGTCTTTGCGTTTCTTTTATATGTTGCCACCTTTATGTATGTATTTT
CTACGTTTGCTAACATACTGCGTAATAAGGAGTCTTAATCATGCCAGTTCTTTTGGGTATTCCGTTAT
TATTGCGTTTCCTCGGTTTCCTTCTGGTAACTTTGTTCGGCTATCTGCTTACTTTTCTTAAAAAGGGC
TTCGGTAAGATAGCTATTGCTATTTCATTGTTTCTTGCTCTTATTATTGGGCTTAACTCAATTCTTGT
GGGTTATCTCTCTGATATTAGCGCTCAATTACCCTCTGACTTTGTTCAGGGTGTTCAGTTAATTCTCC
CGTCTAATGCGCTTCCCTGTTTTTATGTTATTCTCTCTGTAAAGGCTGCTATTTTCATTTTTGACGTT
AAACAAAAAATCGTTTCTTATTTGGATTGGGATAAATAATATGGCTGTTTATTTTGTAACTGGCAAAT

TAGGCTCTGGAAAGACGCTCGTTAGCGTTGGTAAGATTCAGGATAAAATTGTAGCTGGGTGCAAAATA
GCAACTAATCTTGATTTAAGGCTTCAAAACCTCCCGCAAGTCGGGAGGTTCGCTAAAACGCCTCGCGT
TCTTAGAATACCGGATAAGCCTTCTATATCTGATTTGCTTGCTATTGGGATTTAAATCGATGAGTGCG
GTACTTGGTTTAATACCCGTTCTTGGAATGATAAGGAAAGACAGCCGATTATTGATTGGTTTCTACAT
GCTCGTAAATTAGGATGGGATATTATTTTTCTTGTTCAGGACTTATCTATTGTTGATAAACAGGCGCG
TTCTGCATTAGCTGAACATGTTGTTTATTGTCGTCGTCTGGACAGAATTACTTTACCTTTTGTCGGTA
CTTTATATTCTCTTATTACTGGCTCGAAAATGCCTCTGCCTAAATTACATGTTGGCGTTGTTAAATAT
GGCGATTCTCAATTAAGCCCTACTGTTGAGCGTTGGCTTTATACTGGTAAGAATTTGTATAACGCATA
TGATACTAAACAGGCTTTTTCTAGTAATTATGATTCCGGTGTTTATTCTTATTTAACGCCTTATTTAT
CACACGGTCGGTATTTCAAACCATTAAATTTAGGTCAGAAGATGAAATTAACTAAAATATATTTGAAA
AAGTTTTCTCGCGTTCTTTGTCTTGCGATTGGATTTGCATCAGCATTTACATATAGTTATATAACCCA
ACCTAAGCCGGAGGTTAAAAAGGTAGTCTCTCAGACCTATGATTTTGATAAATTCACTATTGACTCTT
CTCAGCGTCTTAATCTAAGCTATCGCTATGTTTTCAAGGATTCTAAGGGAAAATTAATTAATAGCGAC
GATTTACAGAAGCAAGGTTATTCACTCACATATATTGATTTATGTACTGTTTCCATTAAAAAAGGTAA
TTCAAATGAAATTGTTAAATGTAATTAATTTTGTTTTCTTGATGTTTGTTTCATCATCTTCTTTTGCT
CAGGTAATTGAAATGAATAATTCGCCTCTGCGCGATTTTGTAACTTGGTATTCAAAGCAATCAGGCGA
ATCCGTTATTGTTTCTCCCGATGTAAAAGGTACTGTTACTGTATATTCATCTGACGTTAAACCTGAAA
ATCTACGCAATTTCTTTATTTCTGTTTTACGTGCAAATAATTTTGATATGGTAGGTTCTAACCCTTCC
ATTATTCAGAAGTATAATCCAAACAATCAGGATTATATTGATGAATTGCCATCATCTGATAATCAGGA
ATATGATGATAATTCCGCTCCTTCTGGTGGTTTCTTTGTTCCGCAAAATGATAATGTTACTCAAACTT
TTAAAATTAATAACGTTCGGGCAAAGGATTTAATACGAGTTGTCGAATTGTTTGTAAAGTCTAATACT
TCTAAATCCTCAAATGTATTATCTATTGACGGCTCTAATCTATTAGTTGTTAGTGCTCCTAAAGATAT
TTTAGATAACCTTCCTCAATTCCTTTCAACTGTTGATTTGCCAACTGACCAGATATTGATTGAGGGTT
TGATATTTGAGGTTCAGCAAGGTGATGCTTTAGATTTTTCATTTGCTGCTGGCTCTCAGCGTGGCACT
GTTGCAGGCGGTGTTAATACTGACCGCCTCACCTCTGTTTTATCTTCTGCTGGTGGTTCGTTCGGTAT
TTTTAATGGCGATGTTTTAGGGCTATCAGTTCGCGCATTAAAGACTAATAGCCATTCAAAAATATTGT
CTGTGCCACGTATTCTTACGCTTTCAGGTCAGAAGGGTTCTATCTCTGTTGGCCAGAATGTCCCTTTT
ATTACTGGTCGTGTGACTGGTGAATCTGCCAATGTAAATAATCCATTTCAGACGATTGAGCGTCAAAA
TGTAGGTATTTCCATGAGCGTTTTTCCTGTTGCAATGGCTGGCGGTAATATTGTTCTGGATATTACCA
GCAAGGCCGATAGTTTGAGTTCTTCTACTCAGGCAAGTGATGTTATTACTAATCAAAGAAGTATTGCT
ACAACGGTTAATTTGCGTGATGGACAGACTCTTTTACTCGGTGGCCTCACTGATTATAAAAACACTTC
TCAGGATTCTGGCGTACCGTTCCTGTCTAAAATCCCTTTAATCGGCCTCCTGTTTAGCTCCCGCTCTG
ATTCTAACGAGGAAAGCACGTTATACGTGCTCGTCAAAGC

-3′

The forward primer for the assembled product:

5′-CGCAAAAGCGGCCTTTAAC-3′

The reverse primer for the assembled product:

5′-GCTTTGACGAGCACGTATAAC-3′

#### RS-dsDNA related

The sequence for the RS-dsDNA:

SwaI restriction site: ***ATTT/TTTA***

5′-

ATACACCTATTCCGGGCTATACTTATATCAACCCTCTCGACGGCACTTATCCGCCTGGTACTGAGCAA
AACCCCGCTAATCCTAATCCTTCTCTTGAGGAGTCTCAGCCTCTTAATACTTTCATGTTTCAGAATAA
TAGGTTCCGAAATAGGCAGGGGGCATTAACTGTTTATACGGGCACTGTTACTCAAGGCACTGACCCCG
TTAAAACTTATTACCAGTACACTCCTGTATCATCAAAAGCCATGTATGACGCTTACTGGAACGGTAAA
TTCAGAGACTGCGCTTTCCATTCTGGCTTTAATGAGGATTTATTTGTTTGTGAATATCAAGGCCAATC
GTCTGACCTGCCTCAACCTCCTGTCAATGCTGGCGGCGGCTCTGGTGGTGGTTCTGGTGGCGGCTCTG
AGGGTGGTGGCTCTGAGGGTGGCGGTTCTGAGGGTGGCGGCTCTGAGGGAGGCGGTTCCGGTGGTGGC
TCTGGTTCCGGTGATTTTGATTATGAAAAGATGGCAAACGCTAATAAGGGGGCTATGACCGAAAATGC
CGATGAAAACGCGCTACAGTCTGACGCTAAAGGCAAACTTGATTCTGTCGCTACTGATTACGGTGCTG
CTATCGATGGTTTCATTGGTGACGTTTCCGGCCTTGCTAATGGTAATGGTGCTACTGGTGATTTTGCT
GGCTCTAATTCCCAAATGGCTCAAGTCGGTGACGGTGATAATTCACCTTTAATGAATAATTTCCGTCA
ATATTTACCTTCCCTCCCTCAATCGGTTGAATGTCGCCCTTTTGTCTTTGGCGCTGGTAAACCATATG
AATTTTCTATTGATTGTGACAAAATAAACTTATTCCGTGGTGTCTTTGCGTTTCTTTTATATGTTGCC
ACCTTTATGTATGTATTTTCTACGTTTGCTAACATACTGCGTAATAAGGAGTCTTAATCATGCCAGTT
CTTTTGGGTATTCCGTTATTATTGCGTTTCCTCGGTTTCCTTCTGGTAACTTTGTTCGGCTATCTGCT
TACTTTTCTTAAAAAGGGCTTCGGTAAGATAGCTATTGCTATTTCATTGTTTCTTGCTCTTATTATTG
GGCTTAACTCAATTCTTGTGGGTTATCTCTCTGATATTAGCGCTCAATTACCCTCTGACTTTGTTCAG
GGTGTTCAGTTAATTCTCCCGTCTAATGCGCTTCCCTGTTTTTATGTTATTCTCTCTGTAAAGGCTGC
TATTTTCATTTTTGACGTTAAACAAAAAATCGTTTCTTATTTGGATTGGGATAAATAATATGGCTGTT
TATTTTGTAACTGGCAAATTAGGCTCTGGAAAGACGCTCGTTAGCGTTGGTAAGATTCAGGATAAAAT
TGTAGCTGGGTGCAAAATAGCAACTAATCTTGATTTAAGGCTTCAAAACCTCCCGCAAGTCGGGAGGT
TCGCTAAAACGCCTCGCGTTCTTAGAATACCGGATAAGCCTTCTATATCTGATTTGCTTGCTATTGGG
***ATTTAAAT***CGATGAGTGCGGTACTTGGTTTAATACCCGTTCTTGGAATGATAAGGAAAGACAGCCGAT
TATTGATTGGTTTCTACATGCTCGTAAATTAGGATGGGATATTATTTTTCTTGTTCAGGACTTATCTA
TTGTTGATAAACAGGCGCGTTCTGCATTAGCTGAACATGTTGTTTATTGTCGTCGTCTGGACAGAATT
ACTTTACCTTTTGTCGGTACTTTATATTCTCTTATTACTGGCTCGAAAATGCCTCTGCCTAAATTACA
TGTTGGCGTTGTTAAATATGGCGATTCTCAATTAAGCCCTACTGTTGAGCGTTGGCTTTATACTGGTA
AGAATTTGTATAACGCATATGATACTAAACAGGCTTTTTCTAGTAATTATGATTCCGGTGTTTATTCT
TATTTAACGCCTTATTTATCACACGGTCGGTATTTCAAACCATTAAATTTAGGTCAGAAGATGAAATT

AACTAAAATATATTTGAAAAAGTTTTCTCGCGTTCTTTGTCTTGCGATTGGATTTGCATCAGCATTTA
CATATAGTTATATAACCCAACCTAAGCCGGAGGTTAAAAAGGTAGTCTCTCAGACCTATGATTTTGAT
AAATTCACTATTGACTCTTCTCAGCGTCTTAATCTAAGCTATCGCTATGTTTTCAAGGATTCTAAGGG
AAAATTAATTAATAGCGACGATTTACAGAAGCAAGGTTATTCACTCACATATATTGATTTATGTACTG
TTTCCATTAAAAAAGGTAATTCAAATGAAATTGTTAAATGTAATTAATTTTGTTTTCTTGATGTTTGT
TTCATCATCTTCTTTTGCTCAGGTAATTGAAATGAATAATTCGCCTCTGCGCGATTTTGTAACTTGGT
ATTCAAAGCAATCAGGCGAATCCGTTATTGTTTCTCCCGATGTAAAAGGTACTGTTACTGTATATTCA
TCTGACGTTAAACCTGAAAATCTACGCAATTTCTTTATTTCTGTTTTACGTGCAAATAATTTTGATAT
GGTAGGTTCTAACCCTTCCATTATTCAGAAGTATAATCCAAACAATCAGGATTATATTGATGAATTGC
CATCATCTGATAATCAGGAATATGATGATAATTCCGCTCCTTCTGGTGGTTTCTTTGTTCCGCAAAAT
GATAATGTTACTCAAACTTTTAAAATTAATAACGTTCGGGCAAAGGATTTAATACGAGTTGTCGAATT
GTTTGTAAAGTCTAATACTTCTAAATCCTCAAATGTATTATCTATTGACGGCTCTAATCTATTAGTTG
TTAGTGCTCCTAAAGATATTTTAGATAACCTTCCTCAATTCCTTTCAACTGTTGATTTGCCAACTGAC
CAGATATTGATTGAGGGTTTGATATTTGAGGTTCAGCAAGGTGATGCTTTAGATTTTTCATTTGCTGC
TGGCTCTCAGCGTGGCACTGTTGCAGGCGGTGTTAATACTGACCGCCTCACCTCTGTTTTATCTTCTG
CTGGTGGT

-3′

The forward primer for the RS-dsDNA:

5′–ATACACCTATTCCGGGCTATACT-3′

The reverse primer for the RS-dsDNA:

5′–ACCACCAGCAGAAGATAAAACAG-3′

### Enzyme digestion

The RS-dsDNA was digested with the restriction enzyme SwaI (restriction site: 5’-ATTTAAAT-3’) (R0604; NEB) with either Buffer 3.1 (B6003; NEB), Buffer 2.1(B6002; NEB), Buffer 4 (B7004; NEB) or Cutsmart buffer (B6004; NEB). The standard digestion was carried out with 5 units of SwaI, 20 ng of RS-dsDNA and 1X buffer inside a 10 µl reaction volume. After digestion, the materials were purified with the Monarch® PCR & DNA Cleanup Kit. For the time point analysis, the reaction mixture was aliquoted into a PCR tube at various time point (1 to 8 mins) and the tube was immediately inactivated at 65°C inside a water bath.

### Modified buffer digestion

For the RS-dsDNA digestion carried out in standard 0.1M NaCl or the 1M NaCl, two adjusted buffers were made for this. The first buffer, denoted 0.1M NaCl buffer was composed of 50mM Tris-HCl, 10mM $MgCl_2$, 100mM NaCl, and 100 µg/ml of BSA (BS-9808K; BioServ UK Limited) at pH 8.0 (same formula as the commercially available buffer 3.1). The second buffer, denoted 1M NaCl was composed of 50mM Tris-HCl, 10mM $MgCl_2$, 1M NaCl, and 100 μg/ml of BSA at pH 8.0. Both buffers were filtered with 0.22 μm PES syringe filter (SLGP033RS; Millipore Sigma) prior to use. The buffers were mixed directly with the restriction enzyme and RS-dsDNA without further dilution. After digestion, the materials were purified with the Monarch® PCR & DNA Cleanup Kit.

### Cas9 RNP crRNA sequence

The design of the on target crRNA sequence was through the IDTDNA Custom Alt-R™ CRISPR-Cas9 guide RNA design online tool.

On target:

```
--------Target DNA sequence------ PAM
/AltR1/UU CCU UAU CAU UCC AAG AAC GUU UUA GAG CUA UGC U/AltR2/
```

Off target 1:

```
--------Target DNA sequence------ PAM
/AltR1/UU CCU UAU CAU UCC AAG AAU GUU UUA GAG CUA UGC U/AltR2/
```

Off target 2:

```
--------Target DNA sequence------ PAM
/AltR1/UU CCU UAU CAU UCC AAG AGC GUU UUA GAG CUA UGC U/AltR2/
```

Off target 3:

```
--------Target DNA sequence------ PAM
/AltR1/UU CCU UAU CAU UCC AAG GAC GUU UUA GAG CUA UGC U/AltR2/
```

Off target 4:

```
--------Target DNA sequence------ PAM
/AltR1/UU CCU UAU CAU UCC AAU AAC GUU UUA GAG CUA UGC U/AltR2/
```

### Agarose Gel Electrophoresis

For the agarose gel electrophoresis, a 1% agarose gel was used. The agarose was mixed with 1X TAE (J75904.K8; Thermo Scientific), melted in a microwave oven at full power for 45 seconds, and casted with the Mini-Sub Cell GT Systems (Bio-Rad). The samples were prepared with purple loading dye (B7024S; NEB), a total of 50 ng of purified DNA (if not already purified, the sample would be purified with the Monarch® PCR & DNA Cleanup Kit prior to analysis) would be mixed with the loading dye. 50 ng of the GeneRuler 1 kb Plus DNA Ladder (SM1331; Thermo Scientific) was used as the DNA ladder. The gel was run at 50 V for 60 minutes, followed by gel staining with Diamond nucleic acid dye as described above. The gel imaging was carried out with the InGenius LHR Gel Doc System (Syngene).

#### Absorption Spectroscopy

The concentration of the dsDNA was measured via absorption spectroscopy using a NanoDrop™ 2000c Spectrophotometer (Thermo Fisher). The instrument was blanked with 1X TE buffer. 2 µl of the sample was dotted onto the measurement platform, and the measurement was performed using an extinction coefficient of 50 mg/ml for $A_{260} = 1$.

#### Scanning Electron Microscopy

The nanopore dimensions were imaged by Scanning Electron Microscopy (SEM) using a Nova NanoSEM at an accelerating voltage of 15 kV. The imaging was performed by Dr Alexander Kulak.

#### Kernel Density Estimation and probability calculations

The Kernel Density Estimation (KDE) is a non-parametric statistical technique used to estimate the probability density function of a random variable from data [11]. This method involves placing a gaussian kernel at each data point and summing them to create a continuous curve that approximates the underlying distribution. We utilise the KDE method to estimate the current amplitude component of the translocation data. The KDE bandwidth is determined by the Scott's rule of thumb method: $bw = (\frac{4\times\sigma^5}{3\times n})^{0.2}$ , where the bw is the bandwidth, $\sigma$ is the standard deviation of the data and n is the number of data.

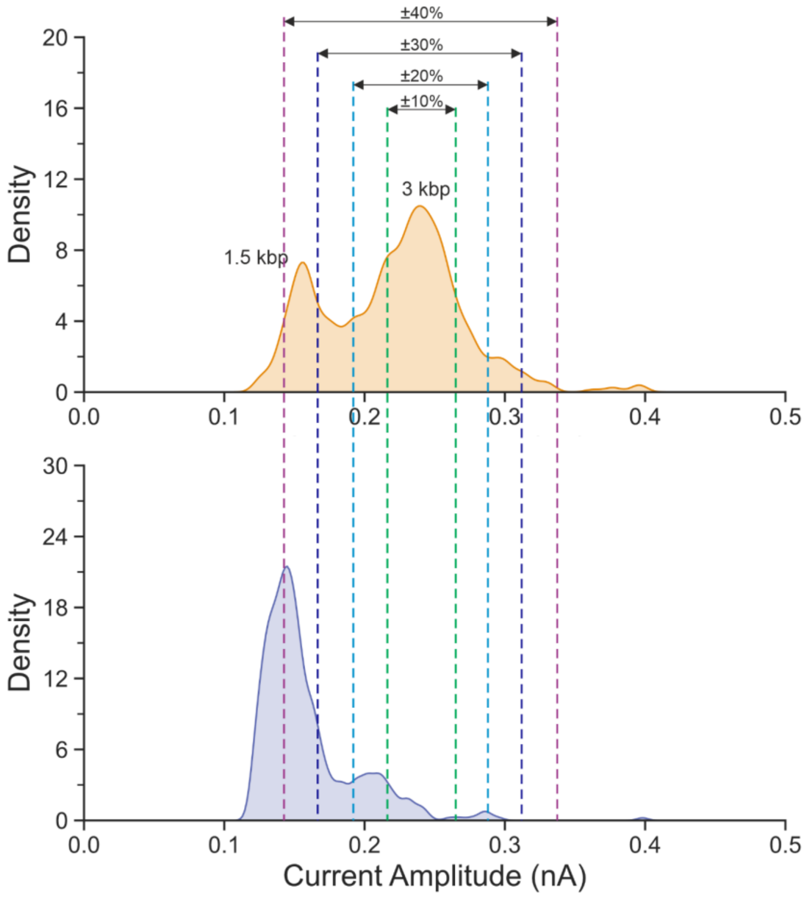

**Supporting Method Figure 1. An example KDE plot of the data from Figure 4.** The data from Figure 4 at 1 min (top0 and 30 minutes (bottom) are shown here. The boundary of either ±10%, ±20%, ±30% or ±40% away from the peak value were shown here.

Using only the 1 min trace as example, which two dsDNA populations exist together: the undigested RS-dsDNA and the digested 1.5 kbp dsDNA, the peak value of the higher conductivity population is identified and two margins of ±10% away from the chosen peak value are calculated. The position of the boundaries is based on filtering of the data, as shown in Supporting Method Figure 1, as the margins move further and further away from the highest point, the boundaries start to include the 1.5 kbp peak at around 0.15 nA. The ±10% is chosen as the margin, it is the most restricted and it increases the specificity, these boundaries are fixed for the rest of the traces (*i.e.* there will be no further determination of the peak value after the first trace). From here, the area under the curve (AUC) within the boundaries of each trace are computed through integration with python.

The total AUC of probability density functions, including KDE, always equate to 1, *i.e.* a 100% probability, the AUC within the boundary thus represents the probability of a translocation event falls into the 3 kbp dsDNA region. These probabilities changes over the reaction time and subsequently, the rate of enzyme reaction can be expressed as: $\frac{\Delta\, AUC\ within\ Boundaries}{\Delta\, Reaction\ Time}$, which can also be phrased as: $\frac{\Delta\, Probabtility\ of\ Detecting\ the\ RS-dsDNA}{\Delta\, Digestion\ Time}$ as seen in Figure 4B.

### Supporting Figure

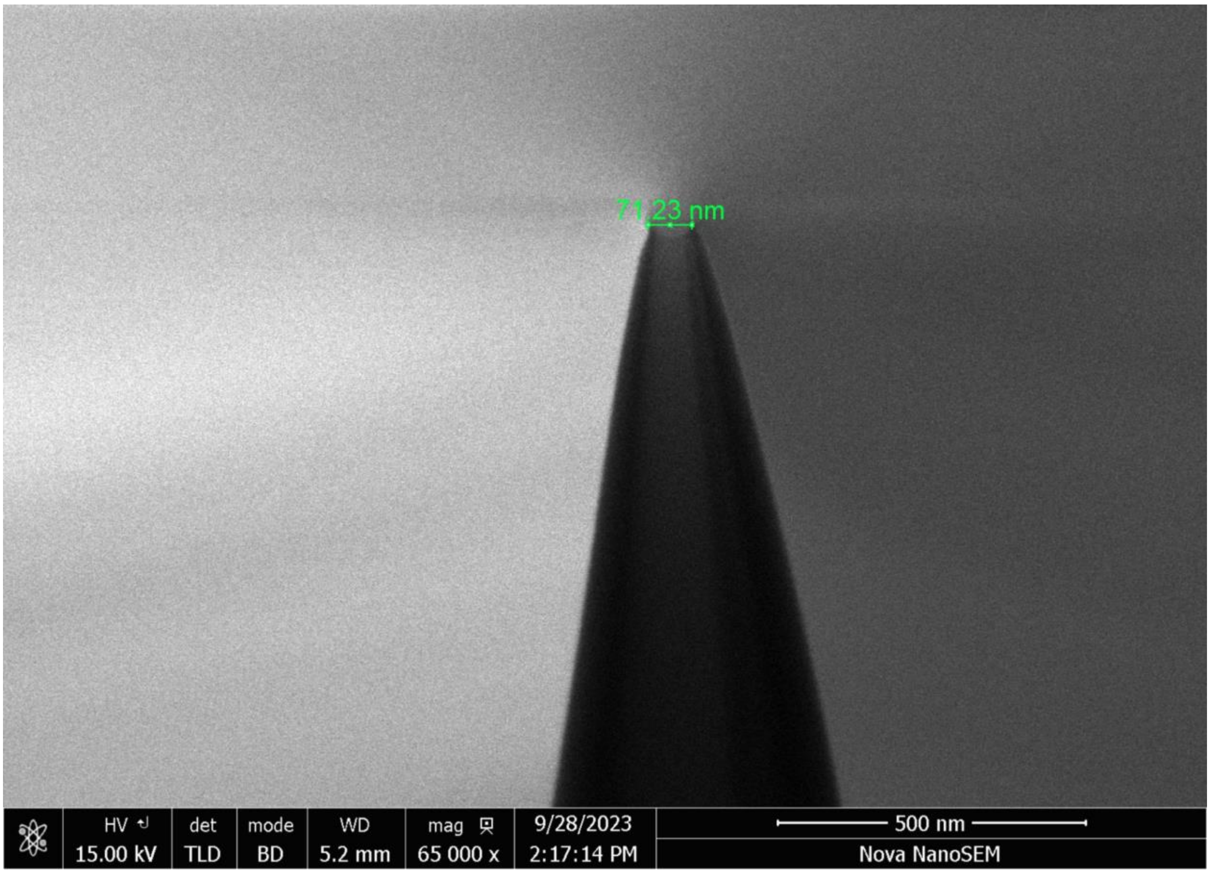

**Supporting Figure 1. Scanning Electron Microscopy micrograph of the nanopipette.** The micrograph shows a nanopore of approximately 70 nm in diameter.

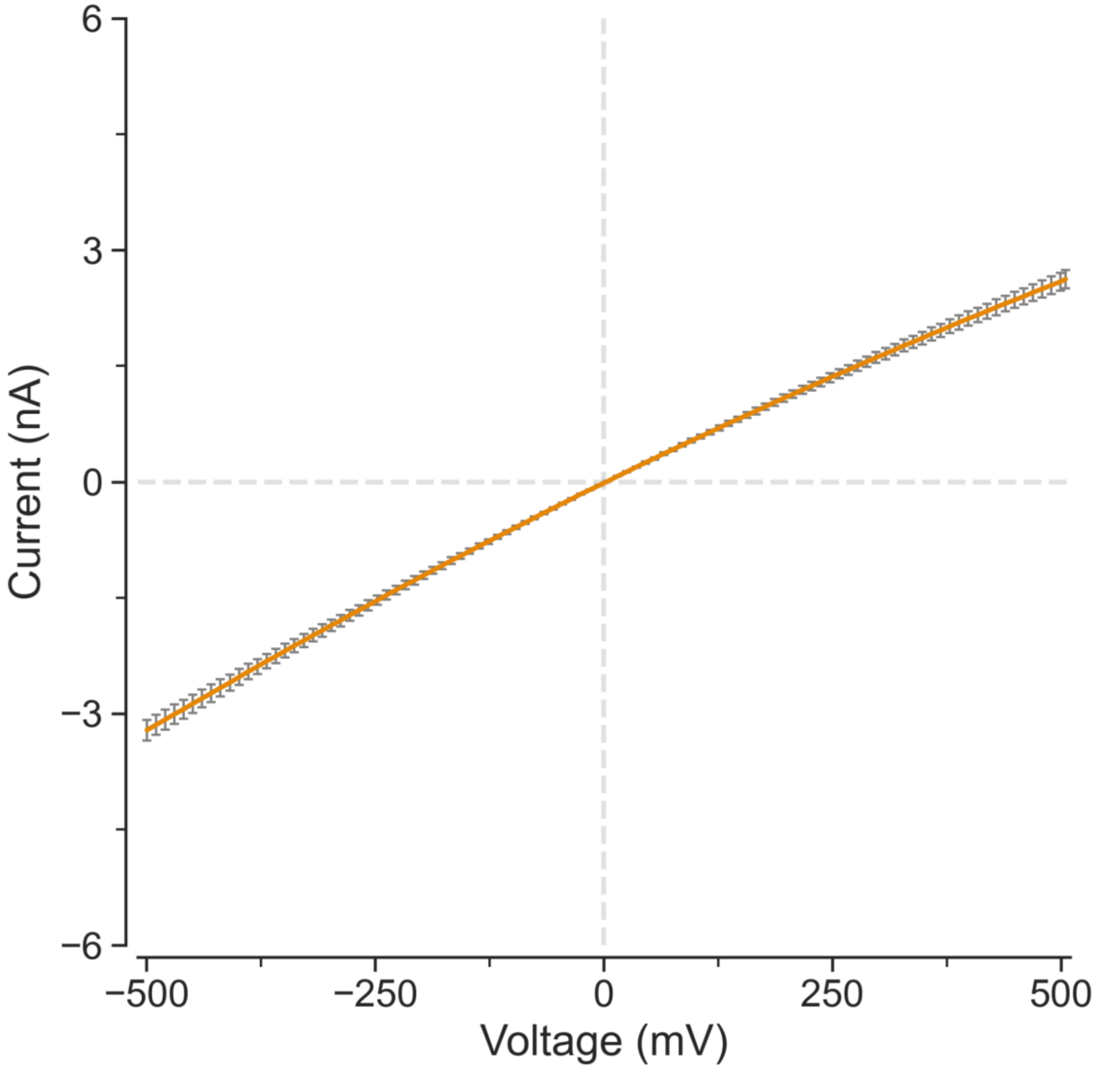

**Supporting Figure 2. The voltammogram of the glass nanopore filled with 0.1M KCl inside a 0.1M KCl electrolyte bath.** The voltammetry measurements of 6 glass nanopore. Error bars are Standard Error of the Mean.

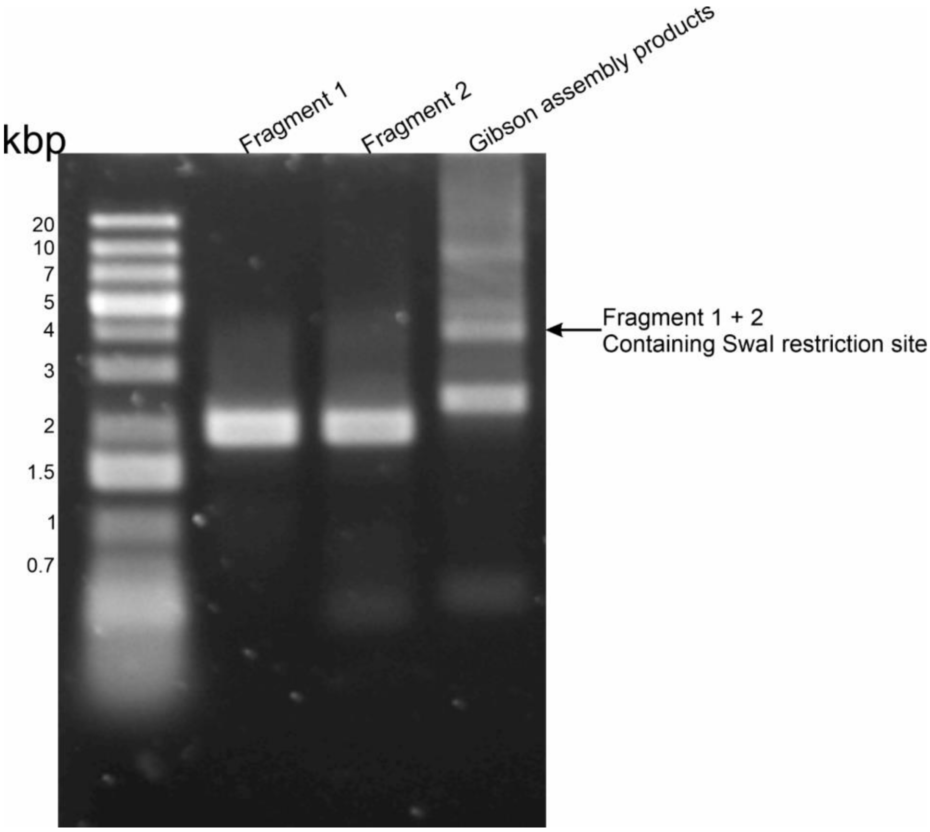

**Supporting Figure 3. Confirmation of the successful assembly between the fragments.** The 4 kbp fragment was isolated from the gel using a blade.

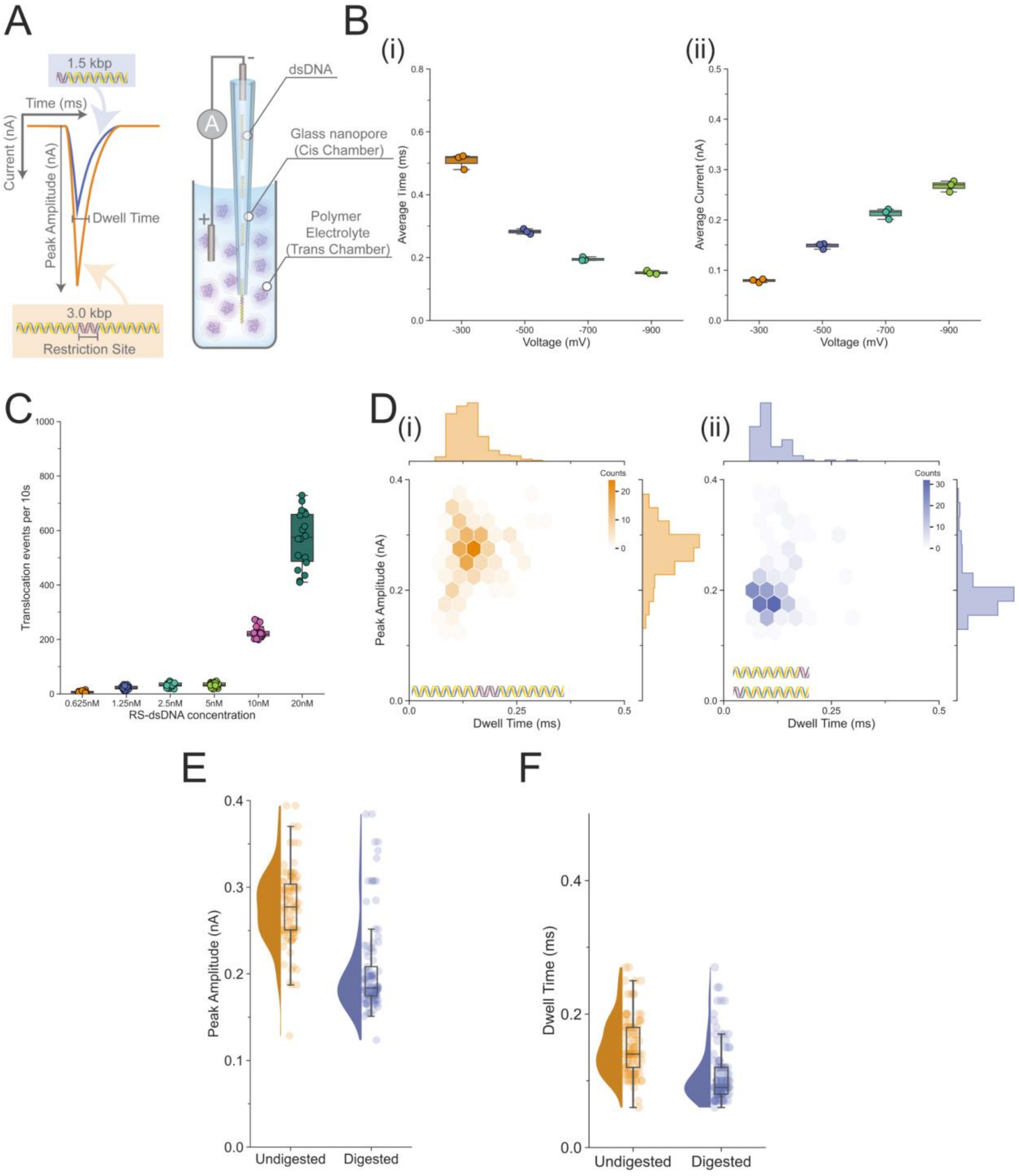

**Supporting Figure 4. Nanopore translocation experiment with the RS-dsDNA. (A)**, Schematic illustration of the translocation signal and the nanopore set-up. The cis chamber contains the analytes (dsDNA) and electrolyte at ~0.1 M monovalent salt, the trans chamber contains a polymer electrolyte blend composed of 0.1 M KCl with 50% (w/v) PEG 35K. The dsDNA is driven from the cis chamber to the trans chamber with a negative voltage and each dsDNA translocation generates a signal with a peak amplitude that depends on the size of the dsDNA. **(B),** The average dwell time (i) and average current amplitude (ii) of the translocation of the RS-dsDNA under -300, -500, -700 and -900 mV. Three separate translocation experiments were carried out. **(C),** Different concentrations of RS-dsDNA ranging from 0.625nM to 20nM were tested. The 10nM was identified as most optimum and used throughout the analysis. **(D)**, The translocation event distribution of the undigested RS-dsDNA (i) and the SwaI digested RS-dsDNA (ii), a population centred at 0.25 ms and 0.15nA could be observed for the RS-dsDNA and a smaller 0.2 ms and 0.10 nA population could be observed for the digested RS-dsDNA. Coloured bars represent the count of events found in each hexagon of the hexbin plot. **(E),** Current peak amplitude distribution of the undigested RS-dsDNA and the digested RS-dsDNA, a clear shift could be seen from 0.3 nA to 0.2 nA. **(F),** Dwell time distribution of the undigested RS-dsDNA and the digested RS-dsDNA. The translocation experiments in **(D-F)** were carried out at -700 mV, the analytes were diluted to 10 nM in 0.1M KCl.

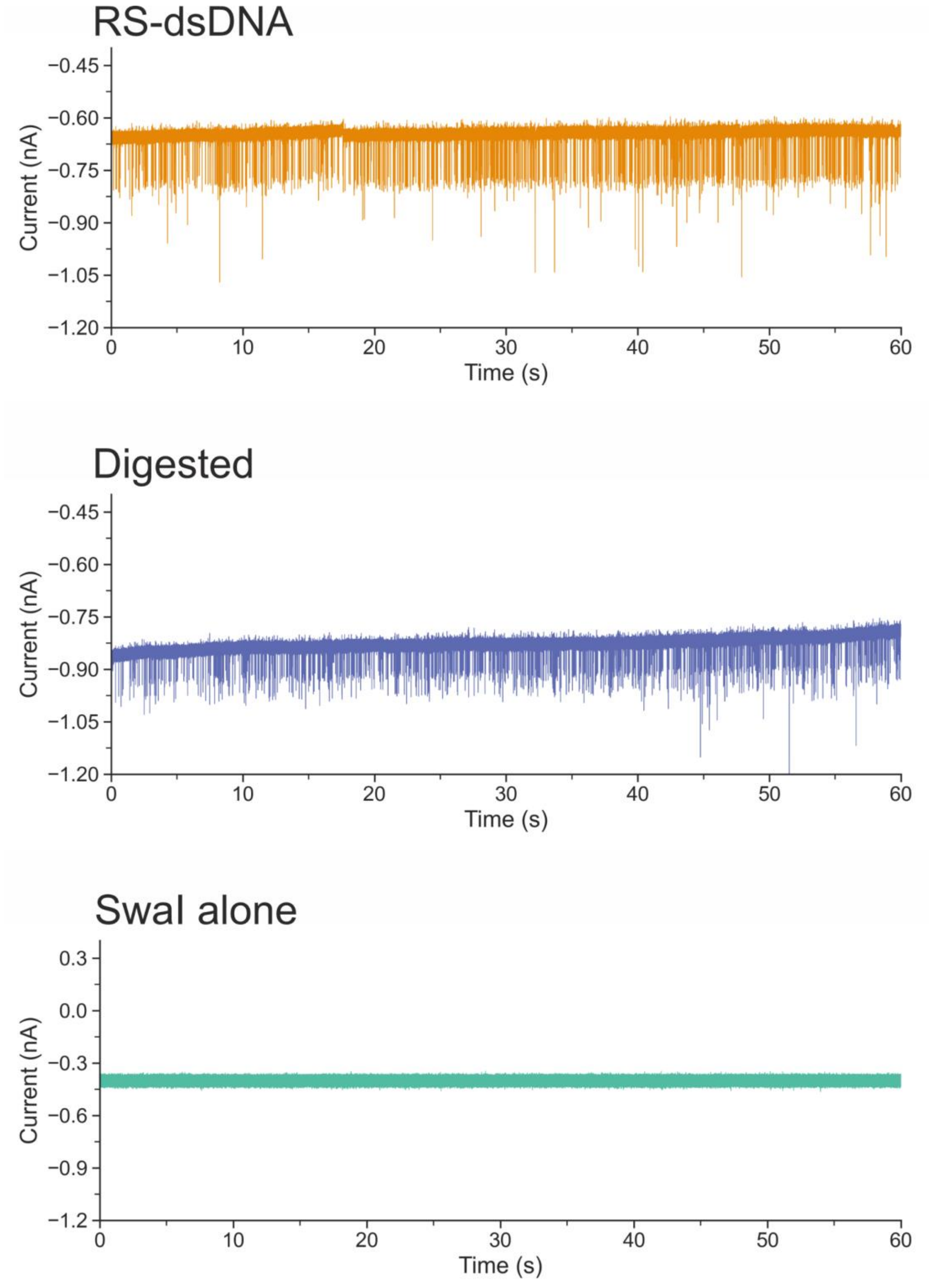

**Supporting Figure 5. The translocation of the RS-dsDNA, SwaI digested RS-dsDNA and SwaI.** The translocation of the RS-dsDNA, SwaI digested RS-dsDNA and SwaI into the polymer electrolyte bath under an applied voltage of -500mV.

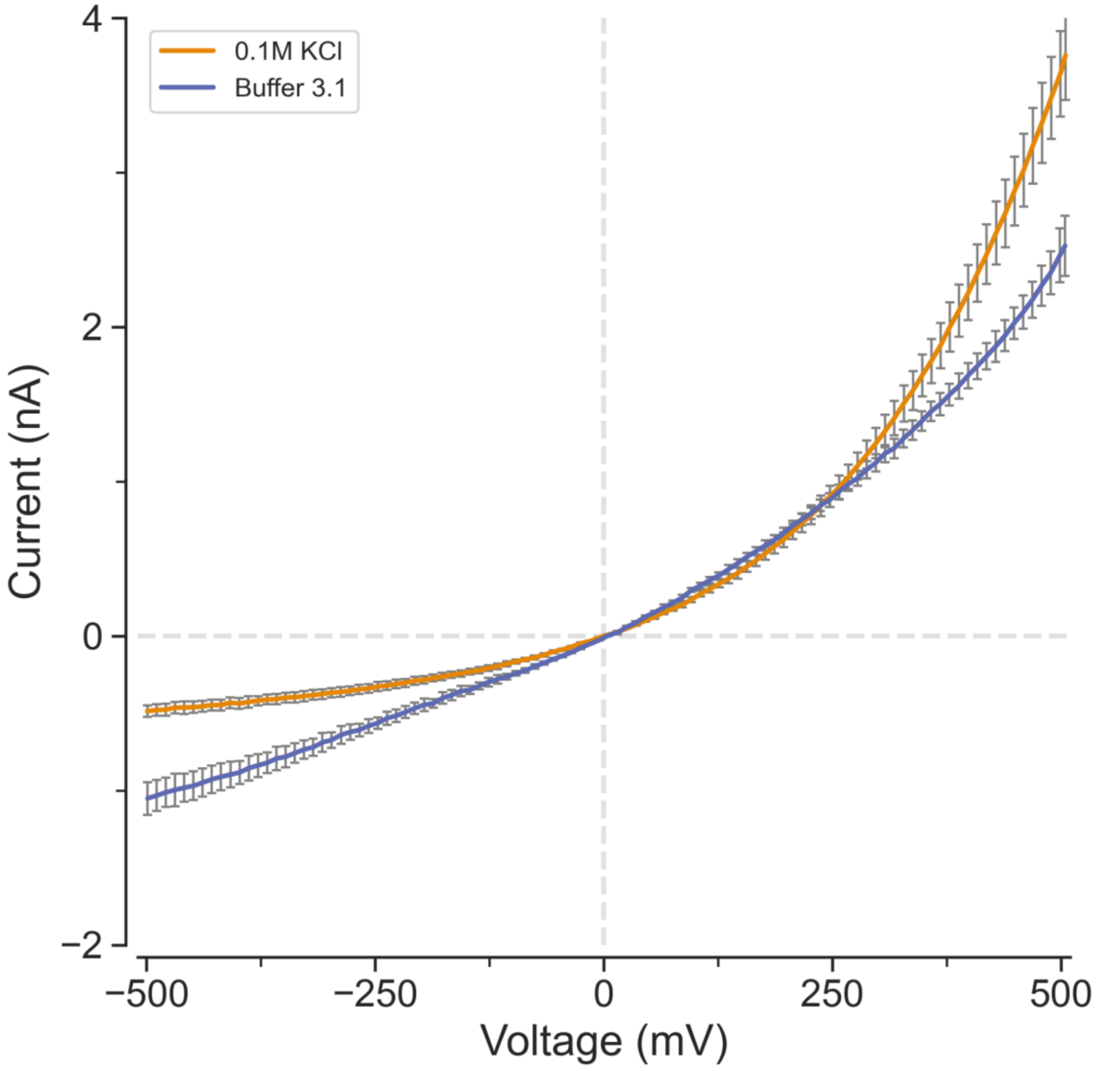

**Supporting Figure 6. The voltammogram of the glass nanopore filled with different buffer inside the polymer electrolyte.** The glass nanopores (3 each for each buffer condition) were filled with either 0.1M KCl or Buffer 3.1 (50mM Tris-HCl, 10mM $MgCl_2$, 100mM NaCl, and 100 µg/ml of BSA at pH 7.9). The significant ion current rectification response was due to the properties of the polymer electrolyte. Both buffers caused the nanopores to be rectified, however, the ICR was more pronounced with 0.1M KCl was used.

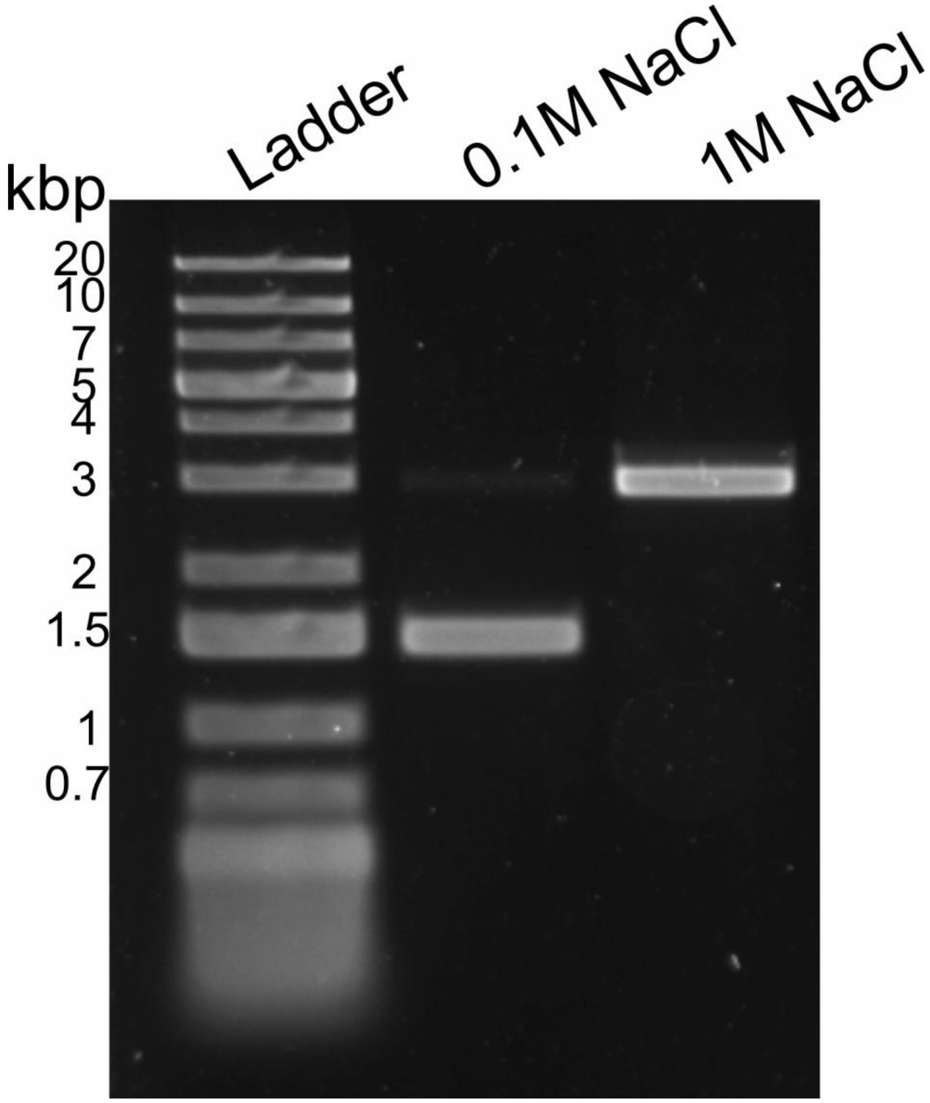

**Supporting Figure 7. Salt concentration dependent digestion of the SwaI enzyme.** Two restriction enzyme buffers were generated, the first buffer contained 50mM Tris-HCl, 10mM $MgCl_2$, 100mM NaCl, and 100 μg/ml of BSA at pH 8.0, this buffer is denoted as the 0.1M NaCl on the gel (This buffer shared the same components as the commercially available one – Buffer 3.1 (New England Biolab) with the exception that the pH was at 8.0 instead of 7.9). The second buffer contained 50mM Tris-HCl, 10mM $MgCl_2$, 1M NaCl, and 100 μg/ml of BSA at pH 8.0, this buffer is denoted as the 1M NaCl buffer. The only difference between these buffers were the concentration of the NaCl. The RS-dsDNA was cleaved into two 1.5kbp dsDNA in the 0.1M NaCl buffer while the 1M NaCl buffer caused the enzyme to become inactive,

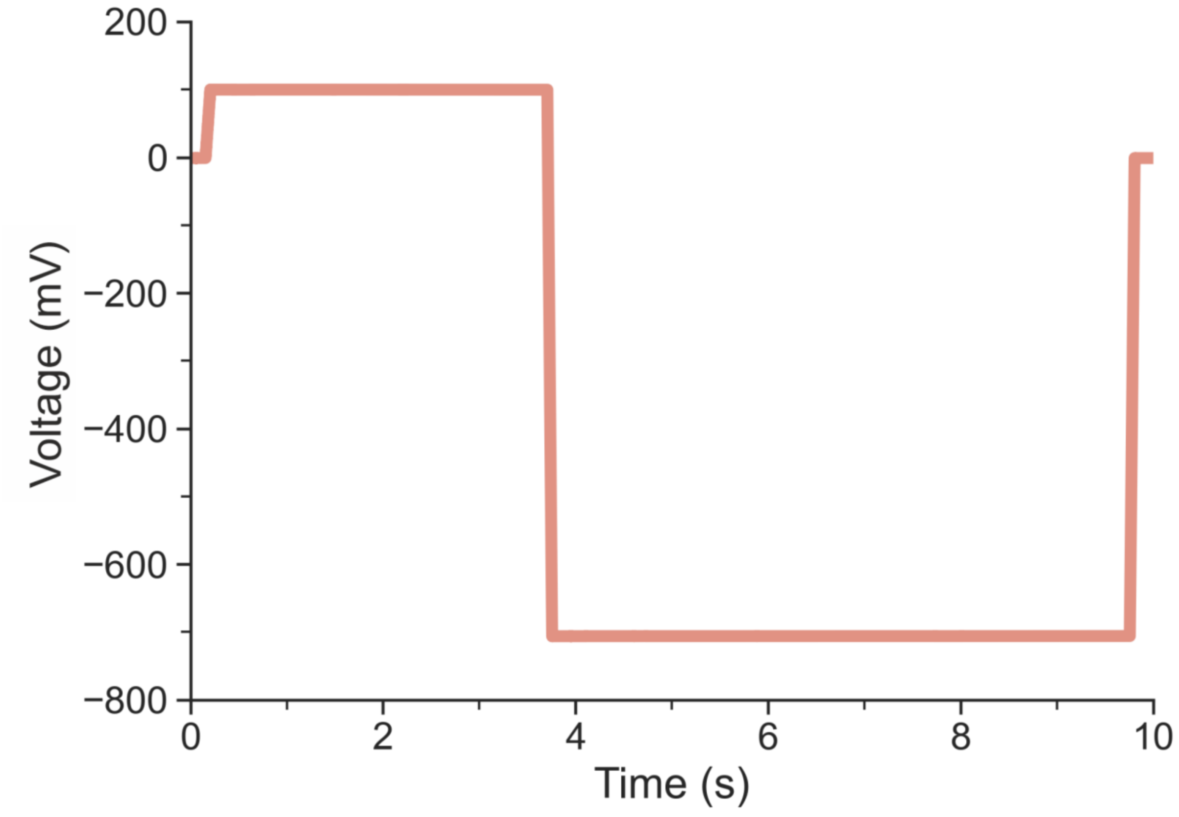

**Supporting Figure 8. The waveform used for the single molecule kinetic measurements.** The voltage first went from 0 to +100 mV for approximately 3.5 seconds, this was followed by a rapid drop to -700 mV for approximately 6 seconds before returning to 0. This waveform would repeat indefinitely until stopped by the user. The waveform was designed to prevent nanopore clogging and control the translocation of the dsDNA.

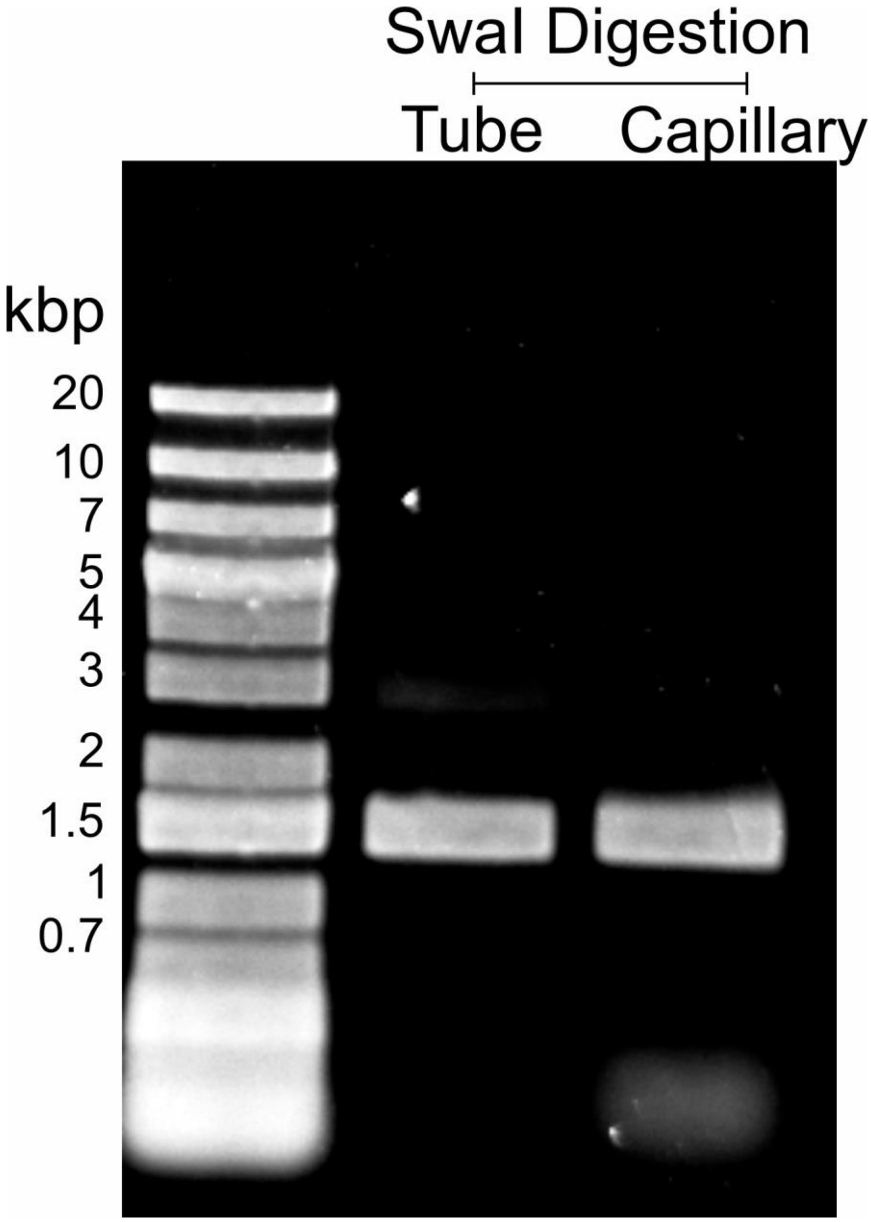

**Supporting Figure 9. RS-dsDNA digestion comparison.** The RS-dsDNA was digested by the SwaI enzyme inside a standard PCR tube or inside a quartz capillary. Both caused the RS-dsDNA to be cleaved into 1.5 kbp dsDNA. Additional small oligos were observed in the capillary digested RS-dsDNA, this could be due to inherent nuclease contamination of the capillary and during handling and extraction of the solution from the capillary.

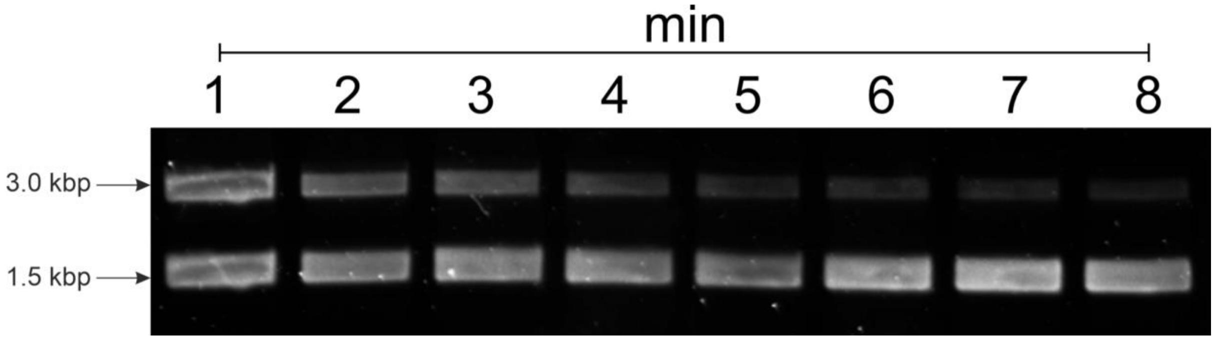

**Supporting Figure 10. The RS-dsDNA digestion kinetic with SwaI.** This gel shows the progression of the digestion from 1 min to 8 mins. The digestion mixture was aliquoted at specific time point into a PCR tube and immediately heat inactivated the restriction enzyme at 65°C. Due to gel well limitation, the ladder was not loaded in this gel.

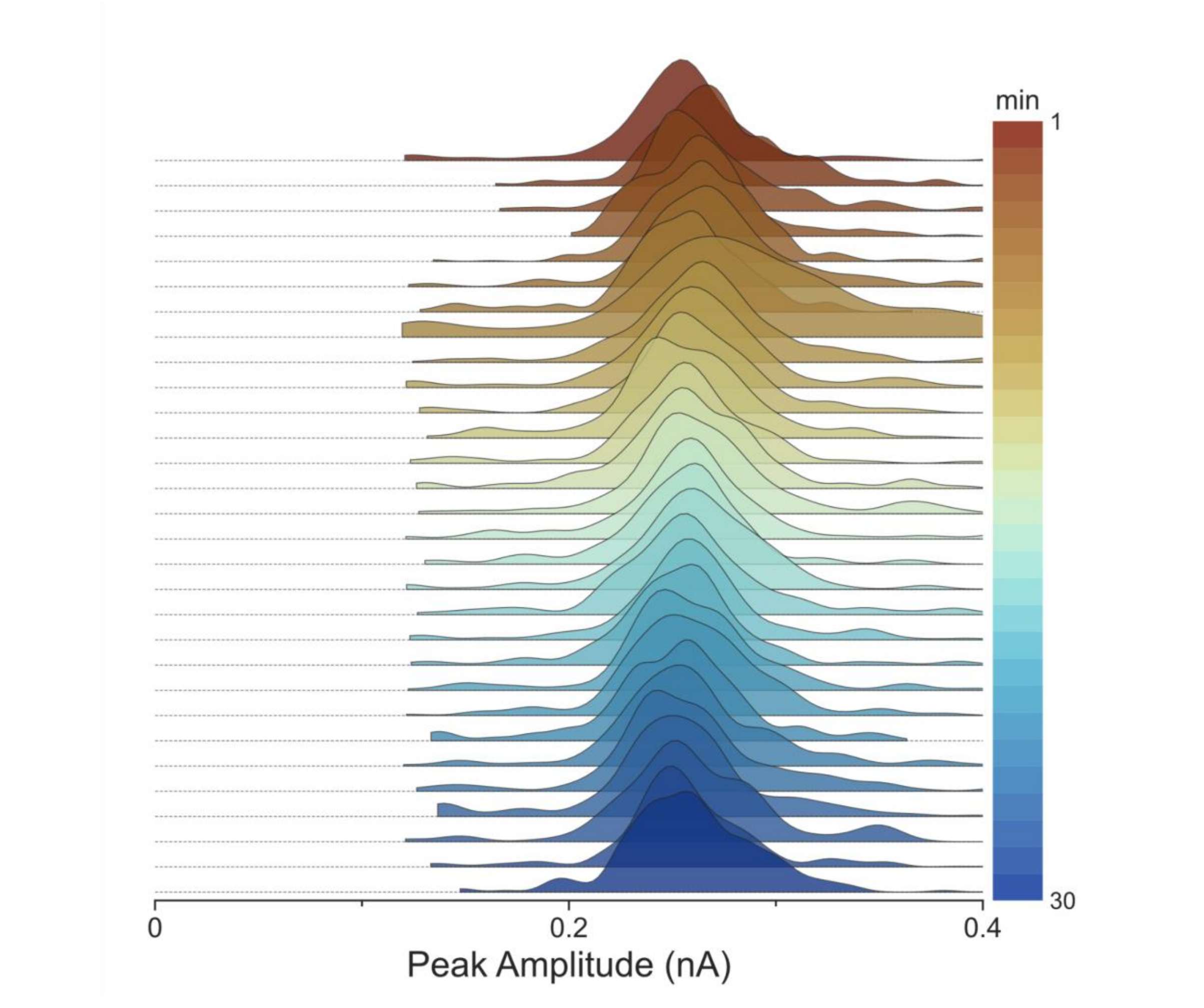

**Supporting Figure 11. Summary of the no enzyme RS-dsDNA translocation data.** The ridgeline plots showing the KDE of all the traces over a period of 30 mins.

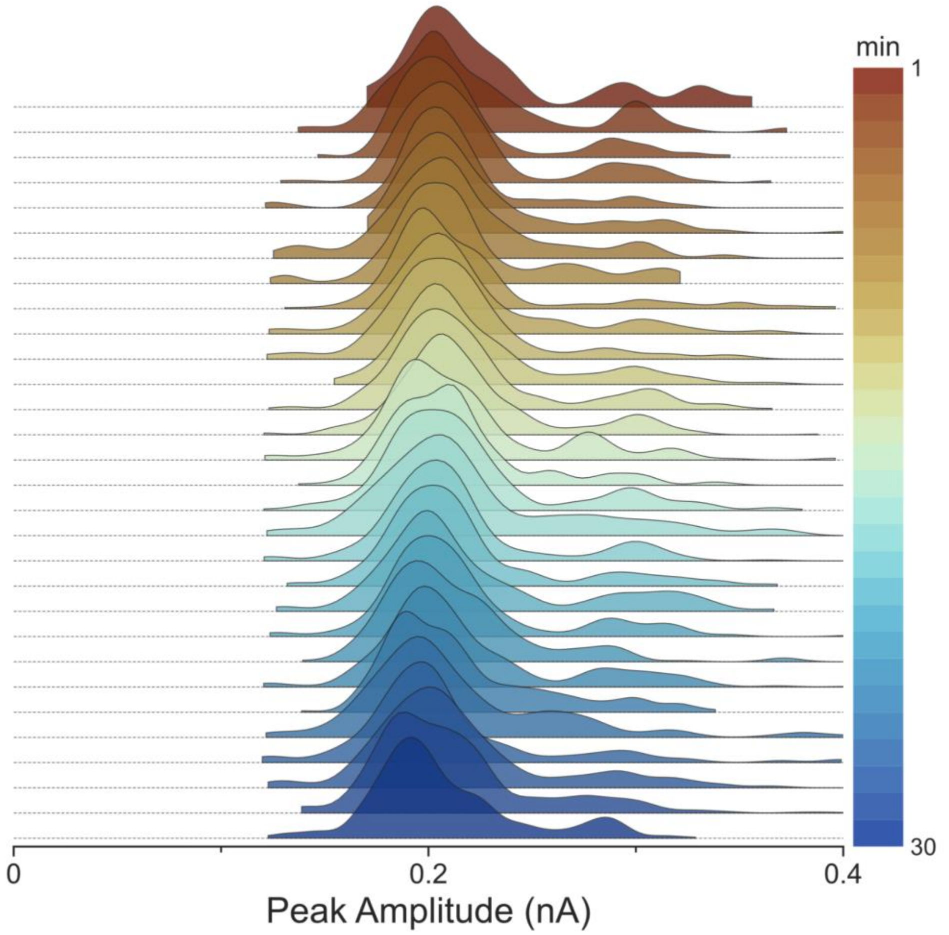

**Supporting Figure 12. Summary of the complete digested RS-dsDNA translocation data.** Top, the ridgeline plots showing the KDE of all the traces over a period of 30 mins.

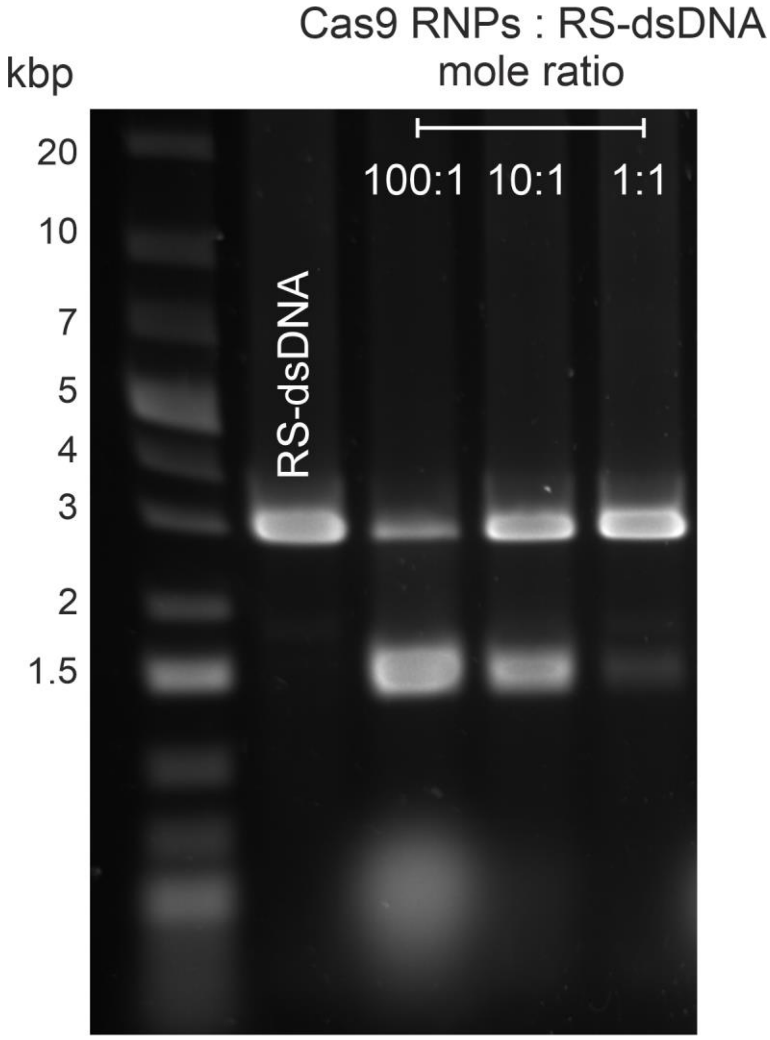

**Supporting Figure 13. Identify the mole ratio of Cas9 RNPs to RS-dsDNA.** Gel electrophoresis was used to analyse the mole ratio between the on target Cas9 RNPs and RS-dsDNA. 100:1 mole ratio was chosen as it leads to the most DNA cleavages.

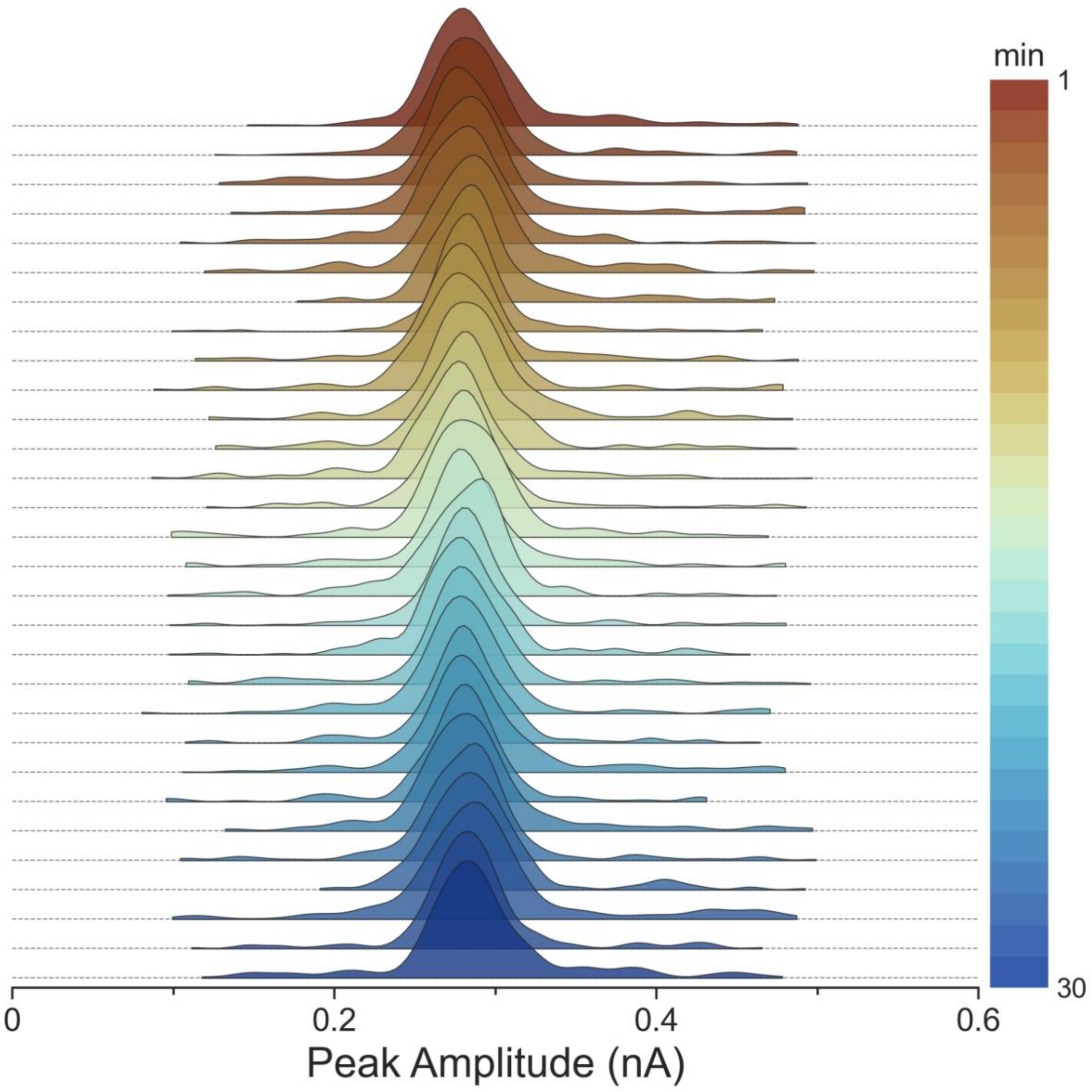

**Supporting Figure 14. Summary of the no Cas9 RNP RS-dsDNA translocation data.** The ridgeline plots showing the KDE of all the traces over a period of 30 mins.

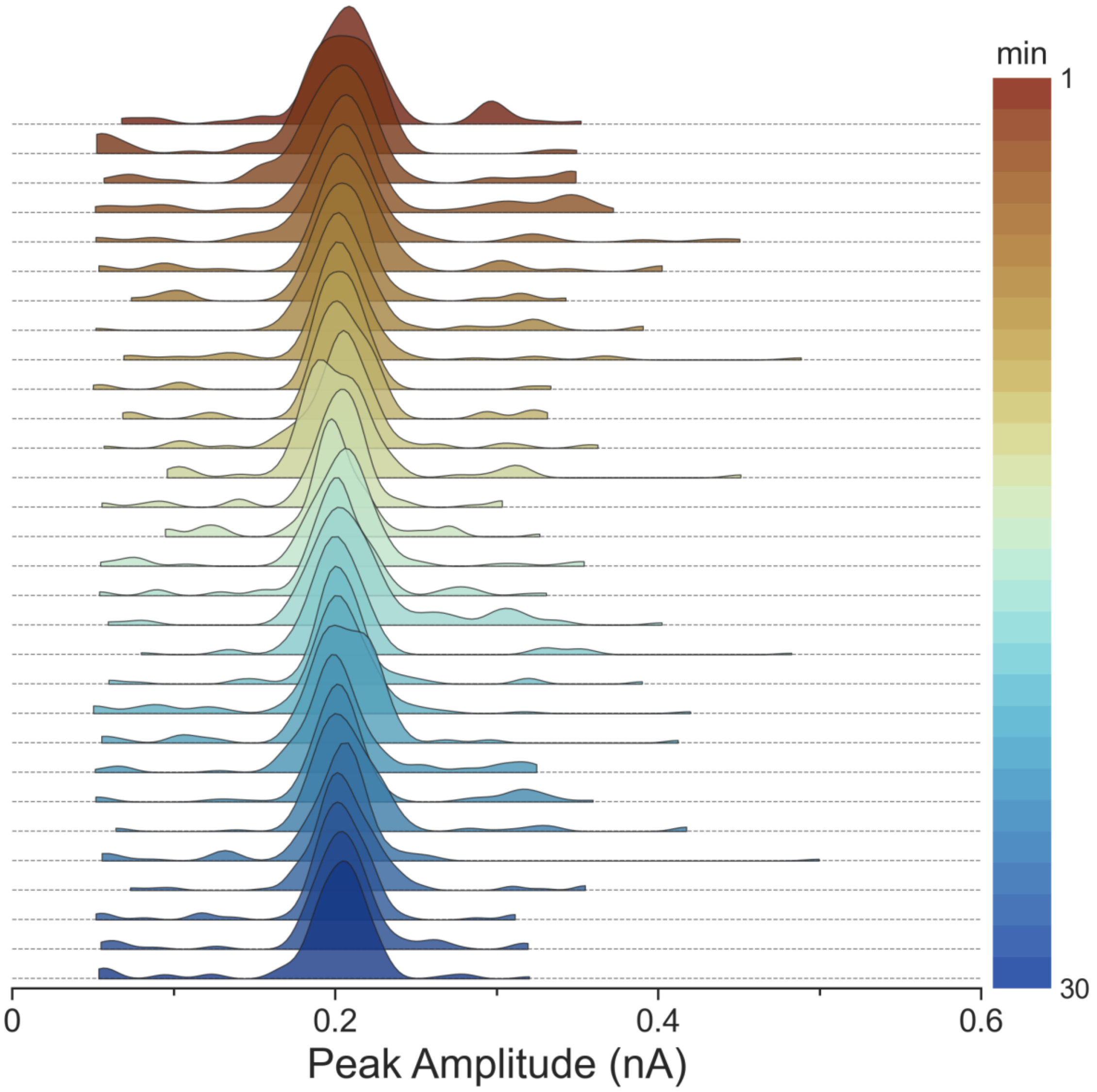

**Supporting Figure 15. Summary of the on target Cas9 RNPs digested RS-dsDNA translocation data.** The RS-dsDNA was pre-incubated with on target Cas9 RNPs for 30 min prior to measurement. The ridgeline plots showing the KDE of all the traces over a period of 30 mins.

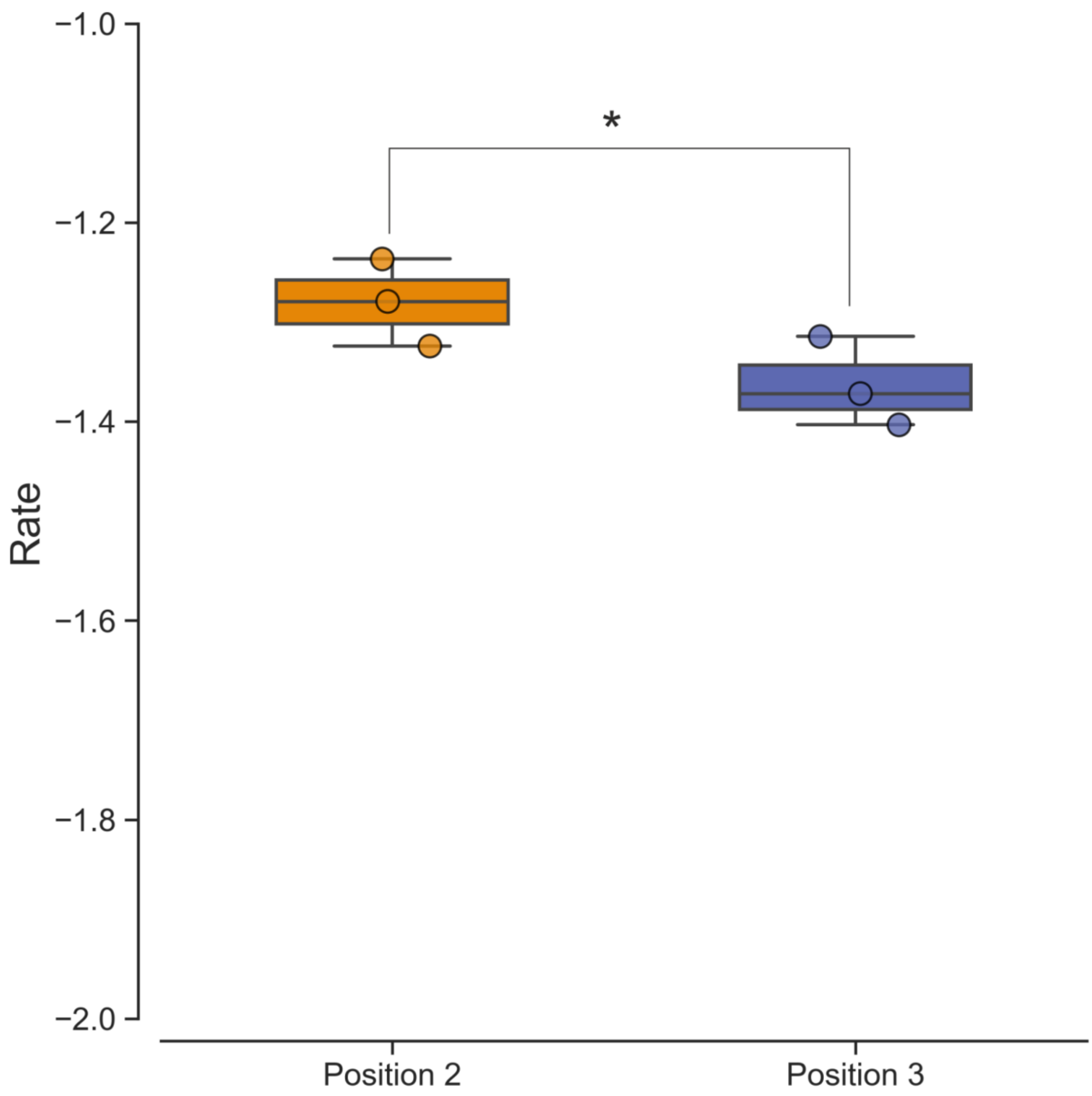

**Supporting Figure 16. Comparison on the reaction rate between off target variant 2 and 3.** Both variants introduced a swap from rA-dT to rG-dT but at either position 2 or position 3 upstream of PAM region. (*, P < 0.05; data assume normal distribution; Levene's test (P>0.05) indicates data are homoscedasticity; one tailed t-test; N = 3 reaction rates).
